# A longitudinal single-cell therapeutic atlas of anti-tumour necrosis factor treatment in inflammatory bowel disease

**DOI:** 10.1101/2023.05.05.539635

**Authors:** Tom Thomas, Charlotte Rich-Griffin, Mathilde Pohin, Matthias Friedrich, Dominik Aschenbrenner, Julia Pakpoor, Ashwin Jainarayanan, Alexandru Voda, Raphael Sanches-Peres, Eloise Nee, Dharshan Sathananthan, Dylan Kotliar, Jason Turner, Saba Nayar, IBD Cohort Investigators, AMP RA investigators, Fan Zhang, Anna Jonsson, Michael Brenner, Soumya Raychaudhuri, Ruth Kulicke, Danielle Ramsdell, Nicolas Stransky, Ray Pagliarini, Piotr Bielecki, Noah Spies, Allon Wagner, Alissa Walsh, Mark Coles, Luke Jostins-Dean, Fiona M. Powrie, Andrew Filer, Simon Travis, Holm H. Uhlig, Calliope A Dendrou, Christopher D Buckley

## Abstract

Precision medicine in immune-mediated inflammatory diseases (IMIDs) requires an understanding of how cellular networks change following therapy. We describe a therapeutic atlas for Crohn’s disease (CD) and ulcerative colitis (UC) following anti-tumour necrosis factor (TNF) therapy. We generated ~1 million single-cell transcriptomes, organised into 109 cell states, from 216 gut biopsies from 38 patients and three controls, revealing disease- and therapy-specific differences. A systems-biology analysis identified distinct spatially-resolved cellular microenvironments: granuloma signatures in CD and interferon (IFN)-response signatures localising to T-cell aggregates and epithelial damage in CD and UC. Longitudinal comparisons demonstrated that disease progression in non-responders associated with myeloid and stromal cell perturbations in CD and increased multi-cellular IFN signalling in UC. IFN signalling was also observed in rheumatoid arthritis (RA) synovium with a lymphoid pathotype. Our therapeutic atlas informs drug positioning across IMIDs, and suggests a rationale for the use of janus kinase (JAK) inhibition following anti-TNF resistance.

## Introduction

Immune-mediated inflammatory diseases (IMIDs) are characterised by impaired immune tolerance leading to chronic inflammation and end-organ damage. The discovery that anti-TNF therapy ameliorates the signs and symptoms of inflammation and tissue damage over three decades ago marked a new era in the treatment of IMIDs^1,2^. However, with non-response rates reaching 40% and the lack of durable remission, medications beyond anti-TNF therapy are required for a large proportion of patients, including many with CD, UC and RA^3–6^.

Recent studies have explored the cellular^7–19^ and molecular basis^20–26^ for these diseases, as well as their associated histopathological features^27^. However, in the gut, the cellular distinctions between inflamed CD and UC, and their respective tissue niches remain poorly understood. Although previous studies have implicated fibroblast activation states^12,13,27^, neutrophils^25–27^, inflammatory monocytes^12,28^, and activated T and IgG^+^ plasma cells^7,12^ with anti-TNF non-response in IBD, no biomarker is currently approved in clinical practice to predict patient response to therapy. Given the absence of validated biomarkers and a plethora of treatment options now available, formulating effective drug sequencing strategies following anti-TNF treatment failure is an unmet clinical need. Understanding the cellular impact of therapeutic agents can inform these strategies. However, no study has directly interrogated the cellular landscape in any IMID before and after anti-TNF therapy using single-cell RNA sequencing (scRNA-seq).

Here, we aimed to create a cell census of two IMIDs, CD and UC to deliver a proof-of-concept therapeutic atlas as a resource for precision medicine. Through the ‘Tissue biomarkers for AdalimUmab in inflammatory bowel disease and RheUmatoid arthritiS’ (TAURUS) study, we sought to understand the implications of gut region and disease activity, as well as the dynamic nature of tissue responses in IBD in the context of the most commonly used class of biologic therapy. Furthermore, we extended our approach beyond the gut to the synovium in RA.

## Results

### A longitudinal scRNA-seq atlas of anti-TNF therapy in CD and UC

We collected biopsies from 38 biologic naïve patients with CD or UC, and three healthy controls across five distinct regions of the gut (terminal ileum, ascending colon, descending colon, sigmoid and the rectum) before and after treatment with adalimumab **(Fig. 1 and Supplementary Table 1)**. Eighty-nine percent of patients (*n*=34) had at least one pair of site-matched biopsies before and after treatment. The mean age of patients with CD and UC was 36 (SD=10.6; range=17-61) and 33 (SD=10.10; range=17-55) years, respectively. Serum trough levels of adalimumab were monitored and patients with anti-drug antibody-mediated loss of response were excluded from the study. Our study comprises 987,743 high-quality single-cell transcriptomes from 216 gut samples **(Fig. 1 and Extended Data Fig. 1a)**. Subclustering of nine cell compartments (myeloid cells, B cells, plasma cells, CD4^+^ T helper cells, CD8^+^ cytotoxic lymphocytes, innate lymphoid cells (ILCs), stromal cells, ileal and colonic epithelial cells) yielded 109 distinct cell states **(Extended Data Fig. 1b, Extended Data Fig. 2, Supplementary Table 2)**.

**Fig. 1.**
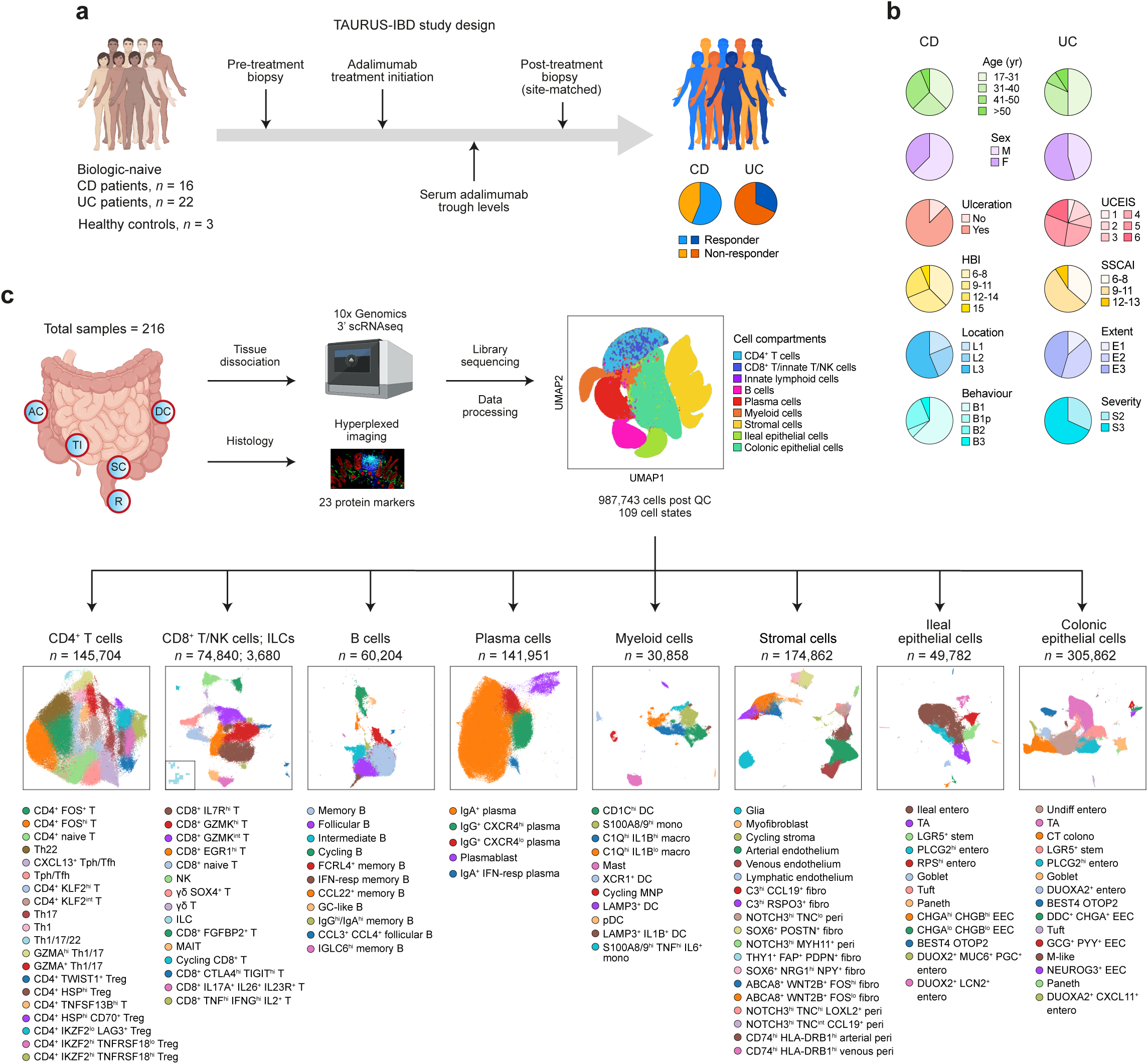
An overview of the TAURUS study. **a,** TAURUS-IBD study design outlining sample collection before and after treatment from biologic naïve patients with IBD. **b,** Clinical characteristics of patients included in TAURUS-IBD. See **Supplementary Table 1** for more details. **c,** TAURUS workflow outlining number of high-quality transcriptomes (987,743 cells) generated across compartments with associated cell states and uniform manifold approximation and projection visualisations. AC, ascending colon; CD, Crohn’s disease; colono, colonocyte; DC, dendritic cell; DC, descending colon; EEC, enteroendocrine cell; entero, enterocyte; F, female; fibro, fibroblast; GC, germinal centre; hi, high; HBI, Harvey-Bradshaw Index; IFN-resp, interferon-responsive; ILC, innate lymphoid cell; lo, low; M, male; macro, macrophage; MAIT, mucosal-associated invariant T; MNP, mononuclear phagocyte; mono, monocyte; NK, natural killer cells; pDC, plasmacytoid dendritic cell; peri, pericyte; R, rectum; RPS^hi^, ribosomal protein S-high; SSCAI, simple clinical colitis activity index; SC, sigmoid colon; TA, transit-amplifying; Tfh, CD4^+^ follicular helper T cell; Tph, CD4^+^ peripheral helper T cell; Th, CD4^+^ T helper cell; TI, terminal ileum; Treg, CD4^+^ regulatory T cell; UCEIS, ulcerative colitis endoscopic index of severity; UC, ulcerative colitis; Undiff, undifferentiated.

### Epithelial heterogeneity drives transcriptomic variation in the ileum compared to colon

Given that variance in our transcriptomic dataset could be attributable to biopsy region or disease type, as well as treatment and associated response, we sought to systematically explore these variables. We first examined healthy samples for differences between terminal ileum and colon across all cell compartments **(Extended Data Fig. 3a-c and Supplementary Table 3)**. Differences were most apparent in the epithelium with 5,493 differentially expressed genes (DEGs) **(Extended Data Fig. 3a)**. Principal component analysis (PCA) demonstrated that 59.7% of variance in the epithelium was explained by the difference between the terminal ileum and colon (PC1) **(Extended Data Fig. 3d)**. PC2 (12.4% of variance) highlighted differences along the colon. Genes involved in the absorption and metabolism of vitamin C *(SLC23A1*), fat soluble vitamins (*RBP2*, *CYP4F2*), vitamin B12 (*TCN2*) and iron (*SLC40A1, CYBRD1*) were preferentially expressed in the terminal ileum, alongside pathways relating to fatty acid metabolism such as unsaturated fatty acid, long-chain fatty acid, as well as triglyceride metabolic processes **(Extended Data Fig. 3e-i**). Distribution of mucin expression varied across ileum and colon. *MUC1*, *MUC4*, *MUC5B*, and *MUC12* were predominantly seen in the colon whilst *MUC17* was preferentially expressed in the terminal ileum **(Extended Data Fig. 3j).**

### A molecular approach to quantifying inflammation across samples

Previous research has highlighted that macroscopically non-inflamed gut samples can nevertheless be histologically and transcriptomically inflamed^13^. Therefore, we generated a gene-based inflammation score using an external dataset examining patient heterogeneity in IBD^27^. We used this gene score to quantify inflammation in our cohort **(Supplementary Table 4, Fig. 2a, and Extended Data Fig. 4a-g)**. Our inflammation score derived from histologically inflamed samples highly correlated with a recently described molecular inflammation score (*R*=0.89, *P* < 2.2×10^−16^) **(Extended Data Fig. 4h)**^29^. The inflammation score was comparable between inflamed samples in both CD and UC **(Fig. 2b)**. We identified several common features in inflamed CD and UC, including cell state expansion in B cells (*CCL22*^+^ memory B), myeloid cells (pDC, S100A8/9^hi^ monocyte and C1Q^hi^ *IL1B*^lo^ macrophage), and stroma (*THY1*^+^ *FAP*^+^ *PDPN*^+^ fibroblast, *NOTCH3*^hi^ *TNC*^hi^ *LOXL2*^+^ pericyte, *NOTCH3*^hi^ *TNC*^int^ *CCL19*^+^ pericyte, and *CD74*^hi^ *HLA-DRB1*^hi^ venous pericyte) **(Fig. 2c-g, Extended Data Fig 4i,j, and Supplementary Table 4)**.

**Fig. 2.**
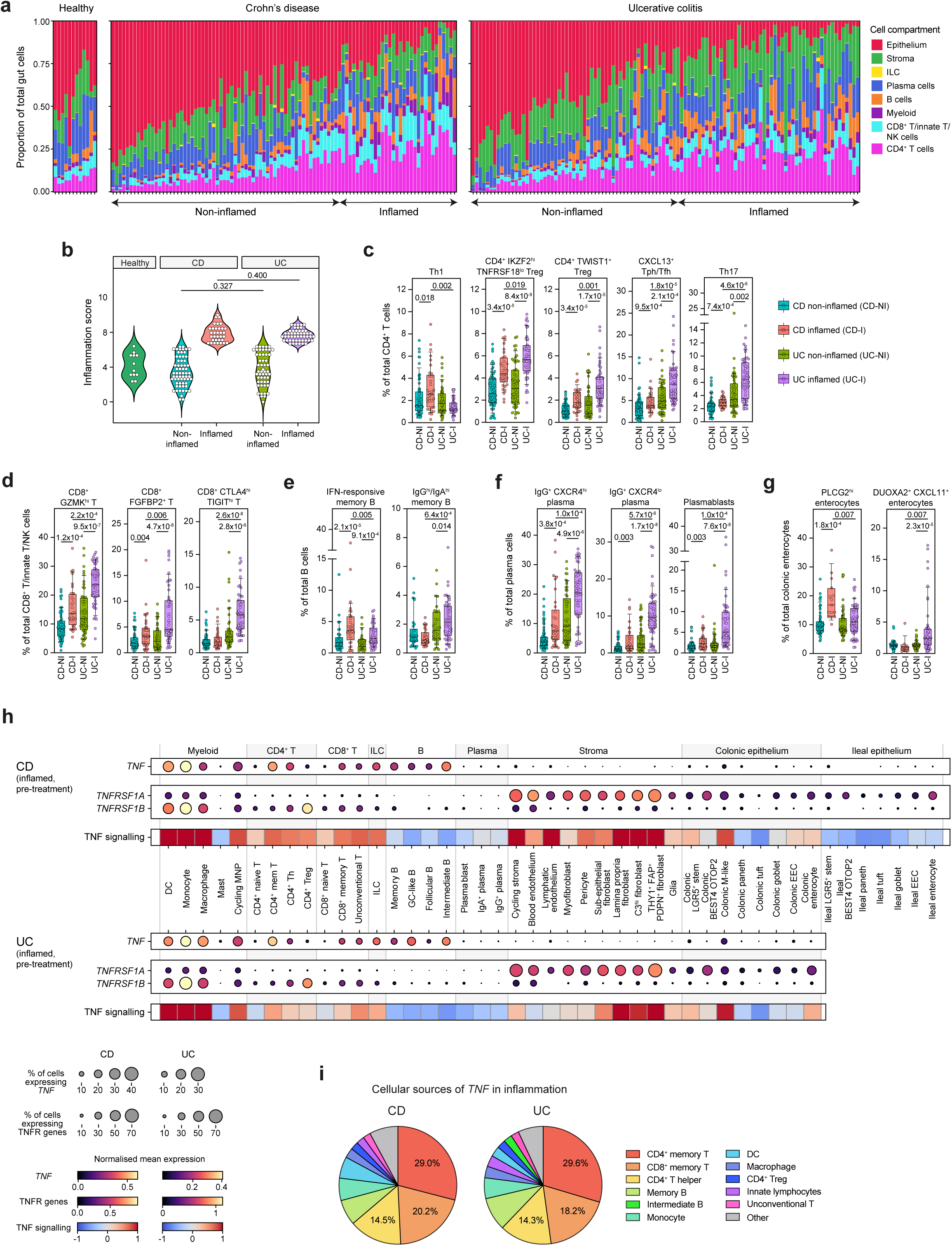
Epithelial and lymphocyte stoichiometry underpins cellular distinctions between CD and UC. **a,** Stacked barplots showing proportion of cell compartments within individual gut samples. Samples are ordered according to inflammatory score. **b,** Violin plots showing distribution of inflammation scores across healthy (*n*=12 samples from 3 patients), CD (*n*=33 inflamed, 63 non-inflamed samples from 16 patients) and UC (*n*=50 inflamed, 53 non-inflamed samples from 22 patients) samples. Wilcoxon rank-sum test used to test significance. **c-g,** Boxplots showing cell state as a proportion of the ‘low’ resolution cell subpopulations (see **Extended Data Fig. 1** for cellular hierarchy), for CD non-inflamed (CD-NI), CD inflamed (CD-I), UC non-inflamed (UC-NI), and UC inflamed (UC-I) gut samples. Sample numbers as in **(b)**. MASC used to test abundance across inflammation status and disease with nested random effects accounting for multiple samples per patient, and covariates including treatment status. Only significant (*P_adj_* < 0.05) differences after multiple comparisons correction with Benjamini-Hochberg are shown. **h,** Mean expression of mRNA transcripts at the ‘intermediate’ cell resolution is shown for *TNF*, *TNFRSF1A* and *TNFRSF1B* in pre-treatment inflamed samples in CD and UC. PROGENy was applied to pre-treatment inflamed samples to calculate TNF signalling scores^38^. Heatmap shows relative enrichment of TNF signalling scores. **i,** Fraction of total TNF transcripts (mean across inflamed samples) at the ‘intermediate’ cell resolution in inflamed samples pre-treatment. DC, dendritic cell; EEC, enteroendocrine cell; entero, enterocyte; GC, germinal centre; hi: high; ILC, innate lymphoid cell; lo, low; MNP, mononuclear phagocyte; Tfh, CD4^+^ follicular helper T cell; Tph, CD4^+^ peripheral helper T cell; Th, CD4^+^ T helper cell; Treg, CD4^+^ regulatory T cell.

### Cellular distinctions between CD and UC are underpinned by differences in lymphocytic and epithelial stoichiometry

Given the distinct clinical and histopathological features of CD and UC, we sought to extract differences between these two conditions at single-cell resolution. In patients with CD, we observed a specific expansion of the Th1 cell state in inflammation **(Fig. 2c)**. Epithelial remodelling in CD consisted of enrichment of an enterocyte cell state (*PLCG2*^hi^ enterocyte), characterised by potent expression of *PLCG2* alongside *PIK3R3* **(Fig. 2g)**. Missense variants of *PLCG2*, which encodes for a phospholipase enzyme, are associated with IBD^30^ and result in intestinal inflammation^31^. Point mutations in the murine orthologue are associated with inflammation driven by autoantibody immune complexes as well as the innate immune system^32^. In addition to being associated with B-cell development^23^ and tuft cells in health^33^, our findings indicate that *PLCG2* expression may be of specific relevance in enterocytes in CD.

Shared features of inflammation were observed in UC and CD, most notably within the B, plasma and CD4^+^ T cell compartments. An IFN-responsive B-cell state (*GBP1*, *STAT1*, *MX1*, *ISG15*) was also found to be more abundant in inflamed samples in CD and UC, although more pronounced in CD **(Fig. 2e and Extended Data Fig. 2c)**. A similar B-cell state has been described in a damage and recovery mouse model of dextran sulfate sodium (DSS) colitis and prevented mucosal healing^34^. Expansion of IgG^+^ plasma cells and plasmablasts were seen in both diseases. However, a greater preponderance of IgG^+^ *CXCR4*^lo^ plasma cell (OR=4.24, 95% CI=2.56-7.02, *P_adj_*=5.45×10^−06^), IgG^+^ *CXCR4*^hi^ plasma cell (OR=2.65, 95% CI=1.75-4.02, *P_adj_*=0.0001), and plasmablasts (OR=2.79, 95% CI=1.79-4.34, *P_adj_*=0.0001) was observed in UC compared to CD **(Fig. 2f)**.

*CXCL13*^+^ Tph/Tfh and Th17 cell states were more abundant in inflamed compared to non-inflamed tissue in both CD and UC **(Fig. 2c)**. However, the expansion of *CXCL13*^+^ Tph/Tfh and Th17 cell states was much more pronounced in UC (OR=2.02, 95% CI=1.58-2.59, *P_adj_*=1.79×10^−05^, and OR=2.27, 95% CI=1.76-2.93, *P_adj_*=4.59×10^−06^, respectively). We observed multiple cell states of CD4^+^ *FOXP3*^+^ regulatory T cells (Tregs)within our dataset **(Extended Data Fig. 2a)**. A select number of these were enriched in UC compared to CD, specifically the CD4^+^ *IKZF2*^hi^ *TNFRSF18*^lo^ Tregs and CD4^+^ *TWIST1*^+^ Tregs (OR=1.26, 95% CI=1.08-1.47, *P_adj_*=0.02 and OR=1.72, 95% CI=1.33-2.24, *P_adj_*=0.001, respectively). *TWIST1* has been reported to be a potent negative regulatory factor which represses Th17 and Tfh, as well as Th1 phenotypes via STAT3 and STAT4 induction, respectively^35,36^.

Given the use of anti-TNF therapy in both CD and UC, we next characterised the expression of *TNF* and its receptors (*TNFRSF1A* and *TNFRSF1B*, that encode TNFR1 and TNFR2, respectively) in our atlas. During inflammation, mean expression of *TNF* on a per cell basis was highest in monocytes and CD4^+^ T memory cells in both CD and UC **(Fig. 2h)**. Of the total *TNF* transcripts detected in inflamed CD and UC, the main cellular sources were CD4^+^ T memory (mean percentage, CD: 29%, UC: 30%), CD8^+^ T memory (CD: 20%, UC: 18%) and CD4^+^ T helper (CD: 15%, UC: 14%) cells **(Fig. 2i).** Although thought to be ubiquitously expressed^37^, *TNFRSF1A* was mainly expressed in structural cells including the epithelium and the stroma, as well as myeloid cells. *TNFRSF1B* was preferentially expressed in immune cells. Beyond profiling cytokine and receptor expression patterns, we also leveraged footprint gene set-based analysis using PROGENy to quantify TNF signalling^38,39^. TNF signalling pre-treatment in inflamed gut samples was relatively higher in CD4^+^ T helper cells, cells of the myeloid lineage, stromal cells, and selected epithelial populations (e.g. M-like cells) compared to B and plasma cells in both CD and UC **(Fig. 2h)**. The relatively diminutive role for TNF signalling in B and plasma cells in IBD is in keeping with the recently suggested association between plasma cells and anti-TNF non-response^7^.

Taken together, this cellular census revealed that CD is characterised by an increase in Th1 cells, as well as expansion of the *PLCG2*^hi^ epithelial cell state. Other changes in the CD4^+^ compartment such as Th17 and *CXCL13*^+^ Tph/Tfh increases, along with IgG^+^ plasma cell expansion, occurred in both diseases but were particularly prominent in UC. However, whilst differences in cell state abundance exist, expression of *TNF*, its receptors, as well as signalling patterns are similar across both forms of inflammatory bowel disease.

### Inflammatory hubs are associated with distinct spatial niches in CD and UC with implications for anti-TNF therapy response

Most scRNA-seq studies rely on partitioning cells into discrete clusters which may not capture the full spectrum of cell identity and activity. We leveraged consensus non-negative matrix factorisation (cNMF) to identify gene expression programmes (GEPs) within cell types^40^. GEPs can represent *bona fide* cell identity but can also be reflective of metabolic processes, activation states as well as response to cytokine signalling occurring concurrently in any individual cell **(Supplementary Table 5 and Extended Data Figs. 5,6)**. We assessed each cell compartment to identify GEPs associated with inflammation **(Supplementary Table 6)** and examined correlations between GEPs within inflamed samples. Groups of correlated GEPs, termed hubs may represent participants in the same or related biological processes. Given the differences between inflamed CD and UC, we derived hubs separately for each disease. This yielded 14 hubs in CD and six in UC **(Extended Data Figs. 7,8)**. Hubs in which more than half of the constituent GEPs were enriched in inflammation were considered to be ‘inflammatory hubs’ **(Fig. 3a,b)**. Correlations were also observed between GEPs from different inflammatory hubs which suggests that these hubs are not mutually exclusive **(Extended Data Fig 7,8)**.

**Fig. 3.**
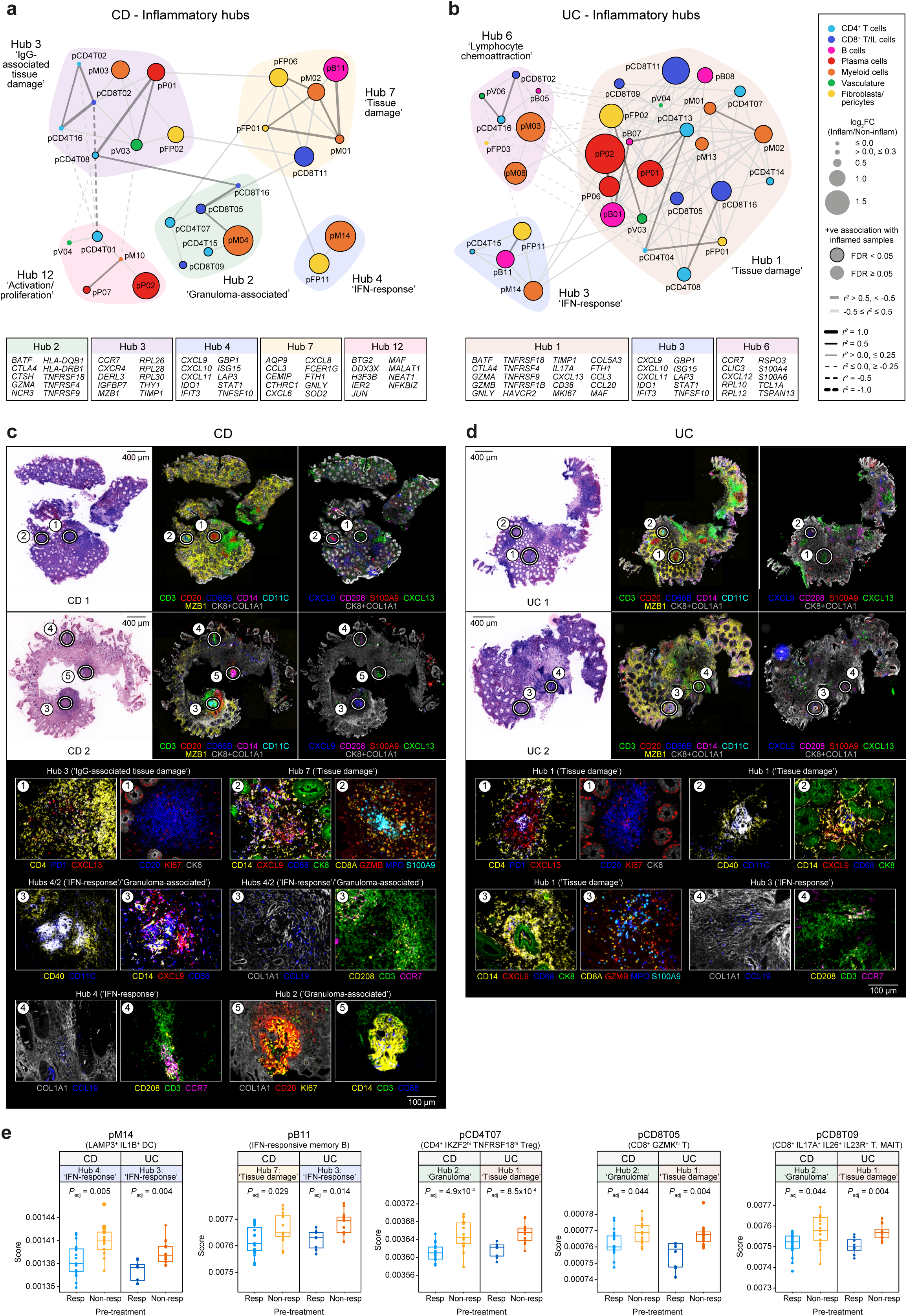
Hubs of gene expression programmes are associated with spatial niches in CD and UC with implications for anti-TNF therapy response. **a,b,** Network graph of covarying GEPs that constitute inflammatory hubs in **(a)** CD and **(b)** UC. Common weighted genes (within top 50) across constituent GEPs within hubs are shown below network graph. See **Supplementary Table 5** for full list of cNMF GEPs in IBD and associated GO term enrichment in GEPs. **c,d,** Virtual H&E with multiplexed imaging highlighting representative regions of tissue and associated protein markers in **(c)** CD and **(d)** UC. Sections shown from two patients from each disease. **e,** Boxplots showing expression of GEPs significantly associated with anti-TNF therapy outcome at baseline in both CD (*n*=36 patients, 17 non-responders, 19 responders) and UC (*n*=24 patients, 16 non-responders, 8 responders) in an external microarray dataset of gut tissue (GSE16879)^23^. Wilcoxon rank-sum test used to assess significance of pre-treatment differences in GEP expression across response status. DC, dendritic cell; FC, fold change; IL, innate lymphoid; MAIT, mucosal-associated invariant T; Non-resp, anti-TNF non-responders; pB, B cell GEP; pCD4T, CD4^+^ T cell GEP; pCD8T, CD8^+^ T cell/NK GEP; pFP, fibroblast and pericyte GEP; pM, myeloid cell GEP; pP, plasma cell GEP; pV, vascular cell GEP; Resp, anti-TNF responders; Treg, CD4^+^ regulatory T cell.

Notably, in both CD and UC, we observed two hubs characterised by response to IFN signalling: hub 4 and hub 3, respectively **(Fig. 3a,b)**. Top weighted genes shared across constituent GEPs included *CXCL9*, *IFIT2*, *IFIT3*, *ISG15* and *STAT1* resulting in enrichment for terms relating to both type I and type II IFN response as well as janus kinase (JAK)/STAT signalling **(Supplementary Table 5)**. Within these hubs, a myeloid and a fibroblast/pericyte GEP (pM14 and pFP11, respectively) were specifically shared between CD and UC **(Fig. 3a,b)**. pM14 was enriched in *LAMP3*^+^ *IL1B*^+^ DCs and to a lesser extent, S100A8/9^hi^ *TNF*^hi^ *IL6*^+^ monocytes **(Extended Data Fig. 6f)**. pFP11 included the follicular reticular cell marker, *CCL19*, trafficking molecules such as *MADCAM1,* selectins *(SELE, SELP*) and MHC class II. Enrichment of this GEP was observed not only in *C3*^hi^ *CCL19*^+^ fibroblasts, but also *CD74*^hi^ *HLA-DRB1*^hi^ venous pericytes in CD and UC **(Extended Data Fig. 6g)**.

We used the CCL19 (pFP11) and CXCL9 (pM14 and pFP11) protein markers to localise the shared GEPs spatially within matched biopsy sections. CCL19 was expressed on COL1A1^+^ stromal cells (pFP11) and also on LAMP3^+^ CCR7^+^ DCs present in CD3^+^ T-cell aggregates **(Fig. 3c,d, region 4**). In our scRNA-seq data, this DC cell state was described by pM08 (*LAMP3*, *CCR7*, *CCL19*) **(Extended Data Fig. 5,6f**) and positively associated with inflammation in both CD and UC **(Supplementary Table 5**). CXCL9 was also found in T-cell aggregates and was expressed on CD14^+^ CD40^hi^ CD11c^+^ monocyte-derived DCs **(Fig. 3c, region 3**). CXCL9^+^ monocyte-derived DCs were additionally situated around damaged epithelial crypt cells **(Fig. 3d, region 2**). Given the expression pattern of CXCL9, this suggests that IFN signalling is associated with inflammation and can be found in both T-cell aggregates and/or regions of epithelial damage in both CD and UC.

Shared GEPs were also seen in hub 7 (CD) and hub 1 (UC). These included pCD8T11 (*FGFBP2*, *GZMB*, *FCGR3A*), pM02 (*S100A8, S100A9*), and pFP01 (*MMP1, MMP3, CXCL5*). These GEPs mapped to CD8^+^ *FGFBP2*^+^ T cells, monocytes and *THY1^+^ FAP^+^ PDPN^+^* activated fibroblasts, respectively. *GZMB*, encoding granzyme B, is a marker of CD8^+^ *FGFBP2*^+^ T cells **(Extended Data Fig. 2b)**. The GZMB^+^ CD8A^+^ T cells localised to areas of epithelial (CK8^+^) damage **(Fig. 3c, region 2,** and **Fig. 3d, region 3)**, in close proximity to S100A9^+^ MPO^+^ CD66B^+^ neutrophil aggregates and CXCL9^+^ monocyte-derived DCs. This suggests that in the context of epithelial damage, CD8^+^ *FGFBP2*^+^ T cells, which potently express *IFNG* **(Extended Data Fig. 2b)**, are driving the IFN response in these monocyte-derived DCs. We have previously described a neutrophil-stromal interaction in context of epithelial damage^27^. Here, we have extended our observations by also localising a CD8^+^ T-cell population, in which granzyme B is detected, to these regions.

In both CD and UC, pCD4T08 (*CXCL13*, *PDCD1*, *CXCR5*) strongly correlated with pP01 (*IGHG1*, *IGHG3*) in hubs 3 and 1 respectively **(Supplementary Table 5)**. These GEPs mapped to *CXCL13*^+^ Tph/Tfh and IgG^+^ plasma cells, respectively **(Extended Data Fig. 6a,d)**. CD4^+^ CXCL13^+^ PD1^+^ Tph/Tfh cells spatially co-localised with CD20^+^ B-cell aggregates in CD and UC **(Fig. 3c,d, region 1)**. This co-localisation was also associated with the presence of plasma cells (MZB1^+^) in the lamina propria, a dominant feature in UC **(Fig. 3d)** but was also observable in some CD patients **(Fig. 3c, CD1)**. This is consistent with our earlier findings comparing the abundance analysis of these cell states in inflamed CD and UC **(Fig. 2c,f)**.

Hub 2 in CD also shared multiple GEPs with hub 1 in UC: pCD4T07 (*FOXP3*, *TIGIT*), pCD8T16 (*CAV1*, *CXCL13*), pCD8T05 (*GZMK*, HLA^hi^) and pCD8T09 (*CTLA4*, *MAF*). Notably, two GEPs (pM04, pCD4T15) present in hub 2 in CD were absent in hub 1 in UC. pM04 was most prominently expressed in a resident macrophage cell state (C1Q^hi^ *IL1B*^lo^ macrophage) **(Extended Data Fig. 6f)**. Top genes in pM04 included *CHIL3L1*, *APOC1, CYP27A1, APOE*, *CTSD*, and *CTSK* **(Extended Data Fig. 5)**. GO term enrichment highlighted multiple terms relating to cholesterol homeostasis and lysosomal transport **(Supplementary Table 5)**. These genes were recently described in the context of granulomatous macrophages in sarcoidosis-affected skin^41^. pCD4T15 (*HLA-DRB1, HLA-DRA, IFNG*) mapped to Th1 and Th1/17 cells **(Supplementary Table 5** and **Extended Data Fig. 6a**). These cells have also been implicated in granulomas in sarcoidosis^41^. This is suggestive of hub 2 being representative of granulomas seen specifically in CD (see **Fig. 3c, regions 3** and **5**). In UC, which is not a granulomatous condition, pCD4T15 was instead strongly correlated with pFP11 within the IFN-response hub 3.

Having placed inflammatory hubs and their associated GEPs in a spatial context, we then assessed whether the GEPs that were positively correlated with inflammation were associated with anti-TNF therapy non-response in an independent cohort (GSE16879)^23^. We discerned multiple GEPs associated with anti-TNF resistance at baseline in this IBD cohort **(Supplementary Table 7 and Fig. 3e)**. Notably, both of the constituent GEPs in the IFN-responsive hub 4 in CD (pM14, pFP11) and more than half of the GEPs comprising hub 3 in UC (pM14, pB11, pCD4T15) were associated with anti-TNF non-response at baseline. Additionally, three lymphocyte GEPs including pCD4T07 (*FOXP3, TIGIT, IKZF2, CTLA4*), pCD8T05 (*GZMK,* HLA^hi^), and pCD8T09 (*CTLA4, IL4I1, MAF, LTB*), which map to CD4^+^ Tregs, CD8^+^ *GZMK*^hi^ T cells, and to CD8^+^ *IL17A*^+^ *IL26*^+^ *IL23R*^+^ T and MAIT cells, respectively, were enriched in non-responders across both CD and UC at baseline.

### Profiling cellular and molecular changes following anti-TNF therapy in CD and UC

We next sought to characterise cellular abundance and molecular changes following treatment in non-responders and responders to adalimumab **(Fig. 4a and Supplementary Table 8)**. No baseline differences in cell state abundance were identified distinguishing responders and non-responders in CD or UC. Rather, in anti-TNF responders, reconstitution of the epithelium occurred, with a corresponding reduction in immune cells. However, this reduction in immune cells differed between CD and UC. A significant reduction in B and plasma cells was specific to UC, potentially reflective of the preponderance of these cell types in this disease. In CD, the reduction was primarily seen in myeloid and T cells. Unlike the case for anti-TNF response, no changes in cell abundance were seen in non-responders with CD, however, an increase in myeloid cells was observed in non-responders with UC.

**Fig. 4.**
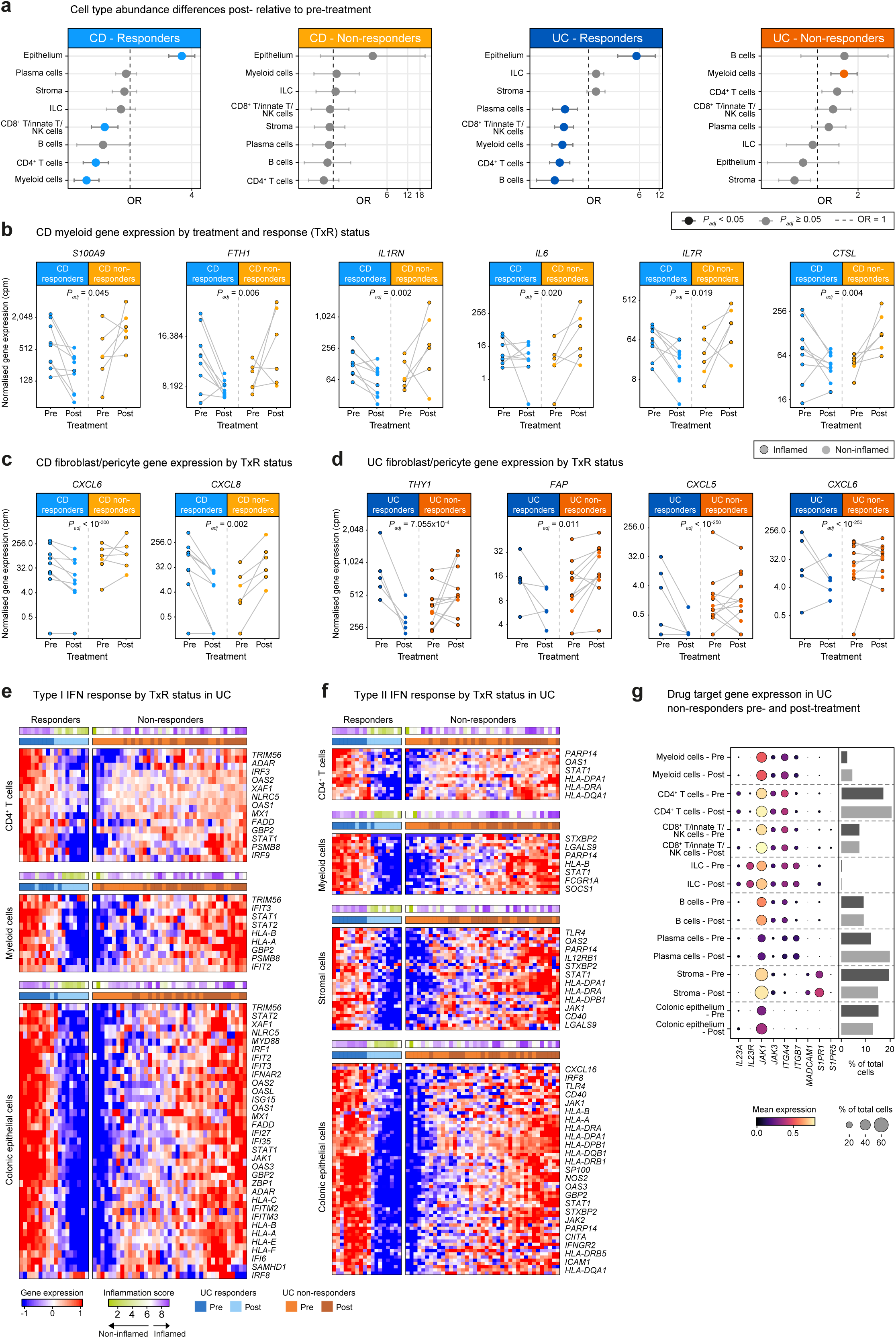
Cellular and molecular changes following anti-TNF therapy in IBD. **a,** Effect of treatment on the cell compartment as a proportion of total cells per sample in anti-TNF responders and non-responders with CD (R: *n*=36 samples from 10 patients, NR: *n*=16 samples from 7 patients) and UC (R: *n*=18 samples from 5 patients, NR: *n*=40 samples from 12 patients). Dots show the odds ratio (OR) and error bars show the 95% confidence interval (CI). **b-d,** Longitudinal changes in gene expression within **(b)** myeloid cells in CD, and **(c)** fibroblasts and pericytes in CD and **(d)** UC. Each paired dot represents median gene expression in a single patient. Significance testing (*P_adj_* < 0.05) performed using *glmmSeq* for treatment:response (TxR) with nested random effects for multiple samples per patient. Only paired samples included. **e,f,** Following over-representation analysis in cell compartments, genes differentially expressed for TxR and associated with type I IFN response (GO:0034340), and type II IFN response (GO:0034341; GO0060333 in CD4^+^ T cells) were examined in the paired samples included in the differential expression analysis. Genes were log-transformed, and scaled from raw counts. The heatmap was generated using *ComplexHeatmap* and split into responders and non-responders with column-wise and row-wise hierarchical clustering. Expression was scaled between −1 to 1. **g,** Dotplot showing expression of genes associated with approved advanced therapies, before and after anti-TNF in non-responders with UC. Bar chart shows median abundance of compartment in context of treatment (pre/post) as a proportion of total cells in sample.

Using an interaction term consisting of treatment and response status^42^, we examined gene expression changes following therapy in CD and UC **(Supplementary Table 9).** In patients with CD who did not benefit from anti-TNF therapy, the largest number of Differentially Expressed Genes (DEGs) was observed in the myeloid compartment. Genes indicative of an inflammatory monocyte phenotype (*S100A9*, *S100A12*, *FTH1*, *IL1RN*) were increased in non-responders but decreased in responders **(Fig. 4b)**. This also included the pro-inflammatory cytokine, *IL6*. GO pathway overrepresentation analysis also highlighted a role for B-cell activation and differentiation in non-responders, in which IL-6 is known to play a role **(Extended Data Fig. 9a and Supplementary Table 10)**. A similar pattern was noted with *IL7R*, which is known to increase in expression on monocytes following exposure to lipopolysaccharide, and is associated with active spondyloarthritis^43^. Inflammatory monocytes also can induce a neutrophil-attractant program in the stroma^27^. Fibroblast and pericyte remodelling following therapy in non-responding patients with CD consisted of neutrophil chemoattractants such as *CXCL6* and *CXCL8* **(Fig. 4c)**.

In non-responders following treatment in UC, markers of the activated fibroblast phenotype (*THY1* and *FAP*), as well as chemokine ligands with potent neutrophil attracting capacity such as *CXCL5* and *CXCL6*, were increased in fibroblasts and pericytes **(Fig. 4d and Extended Data Fig 9b)**. Expansion of THY1^+^ FAP^+^ synovial fibroblasts has been previously associated with RA suggesting that this may be a pathogenic fibroblast cell state in different IMIDs ^16,44^. GO term enrichment revealed persistent IL-1, TNF, and NIK/NF-kappaB signalling pathways alongside neutrophil activation and cell chemotaxis in this compartment despite therapy **(Extended Data Fig. 9b-d and Supplementary Table 10)**. Angiogenesis, as indicated by upregulation of the vascular endothelial growth factor receptor and ephrin receptor signalling pathways, was also active in the stroma **(Extended Data Fig. 9e)**. Evidence of persistent NF-kappaB and TNF signalling was not limited to the stroma but seen across the colonic epithelium, CD4^+^ T cell, B cell, and myeloid compartments **(Extended Data Fig. 10a-d and Supplementary Table 10)**.

Multi-compartmental (CD4^+^ T cell, myeloid, stroma and enterocyte) response to IFN signalling was also seen despite therapy in non-responders with UC. For example, in CD4^+^ T cells, multiple genes indicative of IFN response (*MX1*, *STAT1*, *GBP2*) were upregulated. GO term enrichment analysis suggested response to IFNγ as well as type I IFN **(Fig. 4e,f)**. Plasmacytoid DCs, which are amongst the main producers of type I IFN, were specifically expanded after therapy in non-responders **(Extended Data Fig. 10e, Supplementary Table 8)**. Taken together, these findings suggest that as opposed to stable disease, non-response to anti-TNF therapy is strongly associated with worsening of disease at a cellular and molecular level, indicating a need to promptly switch to alternative therapies in non-responding patients.

Following anti-TNF treatment, a significant reduction in TNF signalling was observed in monocytes, DCs, CD4^+^ T cells, CD8^+^ T cells, and the stromal compartment in responders with CD. In UC, significant decreases were observed in the stromal compartment. Notably in both CD and UC, reductions were also specifically seen in the *THY1*^+^ *PDPN*^+^ *FAP*^+^ activated fibroblast. Changes in TNF signalling in responders negatively correlated with pre-treatment TNF signalling in CD (*R*=-0.43, p=0.0031) and UC (*R*=-0.45, p=0.0049) **(Extended Data Fig. 11a-c)**. This suggests that cells with the largest decrease in TNF signalling following anti-TNF treatment in responders had amongst the highest levels of TNF signalling before treatment.

The ability of anti-TNF to induce mucosal healing represents one of the major advances in the clinical management of IBD. Outcome measures in IBD have now evolved to incorporate histological remission, and most recently, molecular remission^29^. We sought to ascertain whether anti-TNF could also induce cellular remission. We conducted PCA analysis to examine whether non-inflamed post-treatment samples from sites which were inflamed pre-treatment, were distinguishable from non-inflamed pre-treatment or healthy samples. Across the top 10 PCs, there was no correlation between any PC and the treatment variable. Treated samples which achieved cellular remission were comparable to non-inflamed pre-treatment or healthy samples suggesting cellular remission occurs **(Supplementary Table 11)**.

### Mapping targets of advanced therapies in anti-TNF non-response

Finally, we examined targets of approved advanced therapeutic agents for treating IBD across all cell compartments, specifically using paired samples from non-responders to anti-TNF therapy in CD and UC **(Fig. 4g and Extended Data Fig. 12)**. Ustekinumab, vedolizumab, ozanimod, and JAK inhibitors are the options currently available to clinicians to treat patients following failure of anti-TNF therapy. Out of these options, *JAK1* was the only target ubiquitously detected in all cell compartments. Both upadacitinib and filgotinib are particularly attractive given their specificity for *JAK1*. Given the plethora of compounds in development, we also amalgamated a list of cytokines, chemokines, as well as their associated receptors, immune checkpoint molecules, and members of the JAK family to highlight expression at the cell state resolution in the context of genetics and in terms of drug targeting potential to serve as a resource for drug discovery **(Extended Data Fig. 13,14)**. Our resource describes the aforementioned targets specifically using inflamed samples from non-responders to anti-TNF therapy, whilst still on treatment.

### Inflammatory pathways shared between IBD and RA are associated with a lymphoid pathotype in the joint

Shared efficacy to anti-TNF therapy across IMIDs suggests shared biological mechanisms. We therefore wanted to determine whether the cellular hubs and interactions we identified in IBD might underpin inflammation and hold implications for drug response in other anti-TNF responsive diseases such as RA. We recruited patients before and after treatment with adalimumab; *n*=8 patients with paired samples from 4 patients **(Fig. 5a and Supplementary Table 1)**. Whole digestion of synovial tissue following scRNA-seq yielded 65,588 high quality single-cell transcriptomes. We then integrated our data with other whole-digested synovial tissue datasets^15,18^. This resulted in a meta-atlas of 520,603 cells **(Fig. 5a and Extended Data Fig. 15)**.

**Fig. 5.**
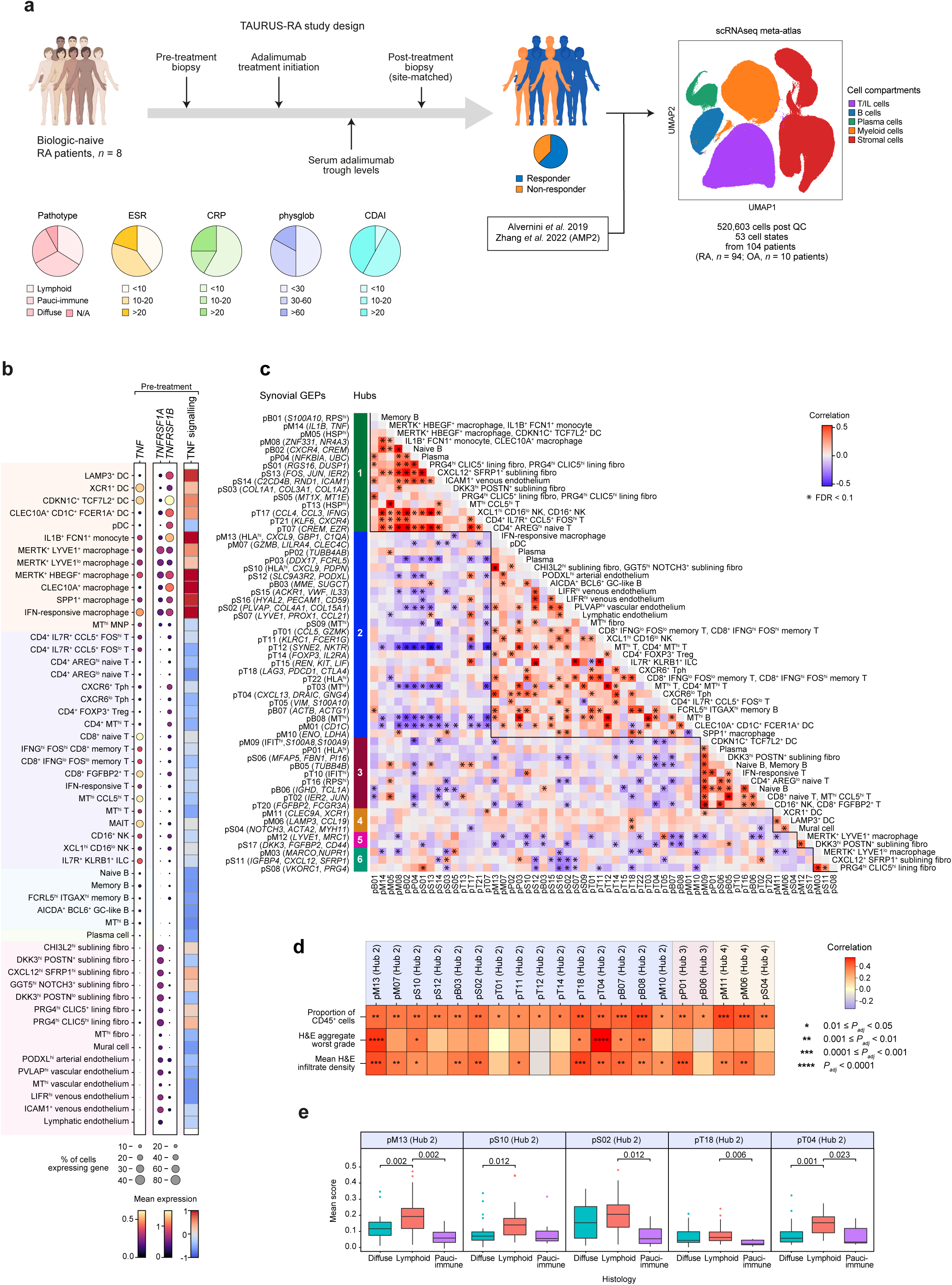
Inflammatory pathways shared between IBD and RA are associated with the lymphoid pathotype in the joint. **a,** TAURUS-RA study design and integration with external datasets to create a meta-atlas for synovial tissue^15,18^. **b,** Mean expression of mRNA transcripts at the cell state resolution is shown for *TNF*, *TNFRSF1A* and *TNFRSF1B* in inflamed samples with RA. PROGENy was applied to inflamed RA samples to calculate TNF signalling scores^38^. Heatmap generated to show relative enrichment of TNF signalling scores. **c,** Correlations of gene expression programmes (GEPs). Asterisk (*) indicates significantly correlated GEP pairs (*P_adj_*< 0.1). Solid lines demarcate hubs of highly correlated GEPs. **d,** Only samples from the AMP2 were included in this analysis as only this dataset had H&E grading for aggregates, and H&E infiltrate density. Spearman correlations between GEP expression and proportion of CD45^+^ cells per sample, worst grade of aggregates and mean infiltration as indicated by associated H&E with FDR correction for number of GEPs within cell compartments. Number of asterisks indicates level of significance. **e,** Associations between GEP expression and histological pathotypes. Only AMP2 data were included in this analysis; diffuse (*n*=30 patients), lymphoid (*n*=33 patients) and pauci-immune (*n*=7 patients) pathotypes. One-way ANOVA conducted to test association between GEPs within cell compartments which were positively correlated with proportion of CD45^+^ cells, with FDR correction for number of GEPs within cell compartments. Pairwise Wilcoxon rank-sum tests only conducted for significant GEPs, with FDR correction for pairwise comparisons between histological pathotypes. Significant adjusted *P*-values displayed above relevant comparisons. CRP, C-reactive protein; CDAI, clinical disease activity index; DC, dendritic cell; ESR, erythrocyte sedimentation rate; fibro, fibroblast; GC, germinal centre; H&E, haematoxylin-eosin; HSP^hi^, heat shock protein-high; IFIT^hi^, Interferon induced proteins with tetratricopeptide repeat genes-high; ILC, innate lymphoid cell; MAIT, mucosal-associated invariant T; MNP, mononuclear phagocyte; MT^hi^, mitochondrial-high; NK, natural killer; OA, osteoarthritis; pB, B cell GEP; pDC, plasmacytoid dendritic cell; physglob, physician global assessment RA; pM, myeloid cell GEP; pP, plasma cell GEP; pS, stromal cell GEP; pT, T/NK cell GEP; RPS^hi^, ribosomal protein S-high; Tph, CD4^+^ peripheral helper T cell; Treg, CD4^+^ regulatory T cell.

Expression of *TNF* was highest in cells of the myeloid lineage as well as T cells **(Fig. 5b)**. Similar to IBD, prominent *TNFRSF1A* expression was seen on stromal cells, whilst *TNFRSF1B* expression was highest on immune cells and specifically myeloid cells. In keeping with our findings from the gut, TNF signalling was highest on myeloid cells and fibroblasts, and relatively lower on B cells and plasma cells in the RA synovium.

Next, we derived cNMF profiles within each cell compartment and associated hubs for RA, as we had done for CD and UC **(Fig. 5c, Extended Data Fig. 16,17, and Supplementary Table 12)**. To determine which GEPs were associated with inflammation, we used a recently developed score for discerning inflammation in the synovium^45^. Twenty out of 58 GEPs across six hubs positively correlated with inflammation **(Fig. 5c,d and Supplementary Table 13)**. We assessed whether GEPs were associated with specific histological features that have been characterised in RA synovial tissue. Fourteen GEPs correlated with infiltrate density, of which 11 belonged to hub 2. Of these five were also associated with aggregates (worst grade) **(Fig. 5d)**. All GEPs enriched in patients with the lymphoid pathotype (pM13, pS10, pT04, pT18) were found in hub 2 **(Fig. 5e)**.

Like hub 4 (CD) and hub 3 (UC) in IBD, genes in multiple GEPs across cell compartments in RA hub 2 (pM13, pS10, and pT22), specifically indicated response to IFN signalling (*GBP1, STAT1, CXCL9*), as well as B-cell activation and proliferation (e.g. *TNFSF13B*) **(Extended Data Fig. 18a)**. pM13 was most enriched in IFN-responsive macrophages, whilst expression of pS10 was most prominent in sublining fibroblast cell states, specifically *CHI3L2*^+^ *GGT5*^+^ *NOTCH3*^+^ and *CXCL12*^+^ *SFRP1*^+^ sublining fibroblasts **(Extended Data Fig. 17)**. Other GEPs such as germinal centre B cells (pB03) and T-cell-associated GEPs facilitating B-cell recruitment (pT04) and activation (pT18) were detected in *CXCR6*^lo^ and *CXCR6*^+^ Tph, respectively, provided further evidence that hub 2 represents a pro-B cell microenvironment.

Given the paucity of well-powered independent longitudinal cohorts examining anti-TNF response using synovial tissue, we sought to examine GEPs in the context of advanced therapy in RA, specifically rituximab and tocilizumab **(Supplementary Table 13**)^42^. Three B cell GEPs (pB03, pB06, pB08), and a plasma cell GEP (pP01) were associated with therapy response to rituximab at baseline **(Extended Data Fig. 18b)**. pB03 and pB06 were indicative of germinal centre (*MME, SUGCT*) and naïve (*IGHD, TCL1A*) B cell states, respectively, whilst pB08 was characterised by mitochondrial (MT-) genes. pB03 and pB08 belonged to hub 2. Although generated from the plasma cell compartment, *MS4A1* (encoding CD20) in addition to multiple MHC class II genes (*HLA-DRA, HLA-DPA1, HLA-DPB1*) were amongst the most top ranked genes in pP01 **(Extended Data Fig. 18c)**. pB06 and pP01 belonged to hub 3. No individual GEP was associated with response to tocilizumab at baseline.

Taken together, these findings suggest that across inflamed gut and joint, there are similarities with respect to *TNF* expression, receptor distribution, as well as cellular responders to TNF signalling. Furthermore, although the constituent GEPs might differ, lymphocyte infiltration programmes associated with IFN signalling are present in multiple cell types across all three IMIDs we studied suggesting that targeting IFN signalling might be considered in these diseases.

## Discussion

Here, we have profiled intestinal tissues at single-cell resolution in CD and UC, before and after administration of the most used biologic agent in the world, adalimumab. This resource represents the first longitudinal, therapeutic scRNA-seq atlas comprising ~1 million cells from 216 samples across 41 individuals (including controls) **(Extended Data Fig. 19)**. This atlas will aid patient stratification and drug discovery efforts in the IBD research community.

We first explored the shared and distinct drivers of inflammation in CD and UC. Although clinically disparate entities, bulk RNA sequencing studies have been limited in their ability to distinguish between them^12^. Through analysis of 145,704 CD4^+^ T cells from the gut, we can confirm Th1 expansion as a hallmark of inflammation in CD but not UC. In addition, we observed a marked expansion of Tph/Tfh cells, IgG^+^ plasma cells, and plasmablasts in UC, as recently reported^7^. However, rather than being characteristic of UC alone, this expansion was observed, albeit to a lesser degree, in CD. Distinctions between both diseases also extended to the epithelium. We describe an enterocyte cell state characterised by *PLCG2* expression that is specifically expanded in inflammation in CD.

Previous studies have endeavoured to uncover patterns of cell abundance that may relate to anti-TNF resistance^12^. However, discrete cell states do not account for continuous phenotypes. Therefore, we supplemented differential cell abundance analysis with cNMF-derived GEPs and identified communities of closely correlated GEPs, termed ‘hubs’ reflective of tissue ecology. Similarities were seen in inflammatory hubs in both diseases such as those characterised by response to IFN signalling, namely hub 4 in CD and hub 3 in UC. Notably, at least half of the constituent GEPs of the IFN-responsive hubs in CD and UC, including pM14 were associated with anti-TNF therapy non-response at baseline.

Using protein markers, pM14 (CXCL9^+^) was present in CD14^+^ CD40^hi^ CD11c^+^ monocyte-derived DCs which localised to two distinct spatial niches: (1) co-occurrence with CCL19^+^ stromal cells (pFP11) in T-cell aggregates and (2) areas of epithelial damage. CCL19^+^ fibroblasts and associated IFN signalling have been described in many IMIDs including RA^45^. We find that this signalling is also present in other stromal cells such as venous pericytes. In UC, pFP11 was strongly correlated with pCD4T15. This GEP is expressed in Th1 and Th1/17 cells which could be the source of IFNγ in this niche. Interestingly, Th1/17 cells have also been implicated in aberrant lymphoid developmental programmes driving granuloma formation in sarcoidosis-affected skin^41^. In CD, we observed that pCD4T15 correlated with pM04. pM04 was characterised by genes considered hallmarks of granuloma-associated macrophages^41^.

In regions of epithelial damage, neutrophil attractant fibroblasts are present^10^. In our data, these cells are represented by pFP01. This GEP was present in the same hub as pCD8T11. pCD8T11 was highly expressed in CD8^+^ *FGFBP2*^+^ T cells, demarcated by *GZMB* expression. Through multiplexed imaging, we observed that GZMB^+^ CD8A^+^ T cells localised to areas of epithelial damage along with S100A9^+^ MPO^+^ CD66B^+^ neutrophil aggregates and the CXCL9^+^ monocyte-derived DCs. As CD8^+^ *FGFBP2*^+^ T cells potently express *IFNG*, they could be driving the IFN response in the DCs within this niche.

Although scRNA-seq studies have previously explored anti-TNF resistance, they have neither directly profiled tissue from responders or non-responders, nor done so in a longitudinal manner^12,13^. As such, the inflammatory landscape following exposure to anti-TNF therapy in non-responders has remained uncharacterised at single-cell resolution. We sought to address this with our longitudinal study design. In CD, we describe an increase in genes associated with inflammatory monocytes (*IL1RN*, *IL6*) in non-responders post treatment. Despite TNF blockade, evidence of persisting and increasing TNF, IL-1 and NF-kappaB signalling was evident in the myeloid compartment in UC. Our group has previously described the relevance of inflammatory monocyte-derived IL-1 and subsequent autocrine signalling in anti-TNF resistance^28^ as well as their capacity to induce a neutrophil attractant programme on fibroblasts^27^. Consistent with these findings, we found an upregulation of genes (*THY1*, *FAP*) indicative of this pathogenic cell state in fibroblasts and pericytes in non-responders with UC.

We also detected evidence of type I and II IFN response increasing across the CD4^+^ T, stroma, myeloid and colonic epithelial cells following treatment in non-responders with UC. Distinguishing between the various IFNs based on transcriptomics alone is challenging. IFNs are pleiotropic cytokines and type I and III IFN can enable epithelial regeneration^46^. The cell-specific, and time-specific role of IFNs, and whether this increase is indeed pathogenic remains unknown. It has been reported that type I IFN released by pDCs contributes to paradoxical psoriasis following the administration of anti-TNF agents^47^ and interestingly, we did see an expansion of pDCs in non-responders with UC. Examining licensed advanced therapeutic agents in the non-responder cohort following treatment demonstrated that *JAK1* was expressed across all cell compartments despite anti-TNF therapy. As such, our data suggest a rationale for why selective modulation of *JAK1* may be effective in the subsequent treatment of anti-TNF non-responders, as appears to be the case in clinical practice ^48–50^.

The amenability of RA to anti-TNF therapy led us to compare across organ systems. We discerned common patterns in terms of *TNF* expression, as well as TNFR distribution in the inflamed gut and synovium. We also detected a hub of GEPs with IFN-response (pM13, pS10, and pT22) enriched in the lymphoid pathotype of patients with RA. As in IBD, we find this IFN-signature is shared with other stromal cell states and with haematopoietic cell states depending on disease context. Thus, we further refine our current understanding of lymphocyte infiltration programmes across IMIDs.

Our longitudinal profiling strategy is a starting point to capture the dynamic evolution of IMIDs at a cellular level over time. To reduce batch effect associated with longitudinal sampling, it was necessary to use frozen samples in this study. This, along with known limitations of droplet-based scRNA-seq did not allow us to detect neutrophils. Another limitation of this study given its observational nature, was the disparity in sampling time after treatment. All patients were sampled after at least eight weeks of exposure to treatment, but sampling time varied up to 1.5 years after therapy initiation due to the COVID-19 pandemic. However, all patients were on therapy at the time of post-treatment sampling.

With the advent of biosimilars and the plethora of available biologic therapies, it is imperative to characterise the impact of individual therapeutic interventions on the cellular landscape in diseased tissues to optimise drug positioning strategies for particular patient subpopulations^51,52^. Therefore, we examined the cellular basis of inflammation and drug response across CD, UC, and RA, specifically in the context of therapy. As the most used first-line biologic, our *in vivo* perturbation atlas of adalimumab serves as a foundation for the investigation of other existing and emerging therapeutic agents in a wide range of IMIDs.

## Methods

No statistical method was used to predetermine sample size, and patients were not randomised as this was an observational study.

### Patient cohorts and ethics

Biologic naïve patients with IBD due to be escalated to adalimumab were recruited from the IBD outpatient clinic at the John Radcliffe Hospital in Oxford. We selected patients with an inflammatory phenotype of CD as biopsies do not always reflect stricturing or penetrating phenotypes affecting the deeper bowel wall layers. Depending on the procedure, biopsies were collected [(IBD Cohort 09/H1204/30)/(GI Ethics 16/YH/0247)] from either terminal ileum, ascending colon, descending colon and rectum (colonoscopy) or the descending colon, sigmoid and rectum (flexible sigmoidoscopy). Clinical history and examination were undertaken to ascertain disease activity; Harvey Bradshaw Index (HBI) for CD and Simple Clinical Colitis Activity Index (SSCAI) for UC. Endoscopic (ulcerative colitis endoscopic index of severity (UCEIS) for UC, and the presence and absence of ulceration for CD) and histologic readouts (Nancy index) were also obtained from the electronic patient records. During follow-up, serum trough adalimumab levels were taken to exclude antibody mediated therapy failure. Samples from the same regions as the pre-treatment endoscopy were taken in post-treatment endoscopy, subject to patient consent and welfare.

Patients with clinically diagnosed RA were recruited to and followed up in an observational standard of care cohort (West Midlands Black Country: 14/WM/1109). Within this, serial synovial biopsies were taken from biologic naïve patients under the following nested ethics (West Midlands Black Country: 07/H1203/57). Patients with RA, according to ACR/EULAR 2010 criteria with a DAS28-ESR score of at least 5.1 and active inflammation in at least one biopsiable joint on ultrasound scanning (GE LogiqE9/6-25MHz probe) underwent ultrasound-guided synovial biopsy. Small joints were biopsied using a spring loaded 16-gauge biopsy needle (Bard Mission); large joints were biopsied using flexible 2.2 mm (Tontarra, Germany) forceps via a single 7 Fr disposable portal placed using Seldinger technique. Four to six synovial fragments were obtained for each small joint needle biopsy, and six to eight synovial fragments from each large joint, taking samples systematically from all available joint recesses. All biopsies were taken by an experienced operator with experience of over 300 procedures; no significant adverse events to biopsy were observed. Clinical assessments (including Disease Activity Score-28) were undertaken at time of biopsy. Patients were re-biopsied in the same joint for follow-up after treatment with adalimumab, subject to patient consent and welfare.

### Obtaining samples and preparation of samples for scRNA-seq

All gut tissue samples were obtained in RPMI 1640 Medium (Gibco) in 50 ml falcon tubes and kept on ice. All samples were processed within 2h of the procedure. Sample processing was performed under sterile conditions. Samples to be used for single-cell analyses were gently washed with 1X PBS, finely macerated with a scalpel and placed into 2 ml of CryoStore CS10 Cell Freezing Medium (CS10; Sigma-Aldrich). Following this, they were kept on ice for 10 min, after which they were transferred to −80°C freezer in Nalgene Mr. Frosty Freezing Containers. After 24 h, they were moved to liquid nitrogen. Samples for histology were placed into formalin for paraffin embedding. Synovial tissue was minced using scalpels to ensure fragments were <1 mm in diameter and randomly assigned into 1 ml cryovials into which 1 ml/vial of CS10 was added. Vials were equilibrated at 4°C for 10 min before transfer into a 4°C Mr. Frosty and storage at −80°C.

### 10X Genomics scRNA-seq library preparation, tissue dissociation and sequencing

Gut tissue samples were thawed into warm IMDM media with foetal bovine serum (FBS). CS10 was removed by washing. Samples were treated with EDTA pre-digestion with rotation for 15 min to remove dead/damaged epithelial cells. Samples were then dissociated enzymatically with Liberase TM and DNase into a single-cell suspension with rotation. Cells were washed, strained and counted for viability using acridine orange/propidium iodide and a maximum of 10,000 cells were loaded per 10X Chromium channel.

Synovial tissue samples were thawed into warm IMDM with 10% FBS and washed two times to remove preservation media. Samples were then digested in a cocktail of Liberase TL and DNase in warm media for 30 min, with agitation. Samples were then strained at 40 microns, and washed two times with PBS with 0.4% BSA. Live events were counted using acridine orange/propidium iodide and a maximum of 10,000 cells were loaded per 10X Chromium channel. The GEX 3’ V3 protocol was followed throughout sequencing for both gut and synovium.

### scRNA-seq pre-processing and quality control filtering

Cell Ranger v3.1.0 was used to align reads to the hg19 human transcriptome and generate feature-barcode matrices from the Chromium single-cell RNA-seq output for TAURUS samples. Panpipes was used to generate anndata objects following quality control, and batch correction^53^. Filtering steps for high-quality single cells included removal of: doublets using Scrublet^54^, cells expressing fewer than 500 genes, and cells with mitochondrial gene count percentage greater than 60%. Genes that were detected in fewer than 3 cells were removed.

### Selection of variable genes, dimensionality reduction, clustering and annotation

To account for differences in sequencing depth across cells, UMI counts were normalised by the total number of UMIs per cell and converted to transcripts-per-10,000. Data were then log-normalised. Highly variable genes were selected following which, a subset of genes consisting of T-cell receptor, immunoglobulin and HLA genes were removed. Data were then scaled prior to PCA. For gut samples, BBKNN was used for batch correction of samples and Leiden clustering was applied to derive broad cell populations^55^. This included: B cells, plasma cells, T cells, myeloid cells, ileal epithelial cells, colonic epithelial cells, and stromal cells. In the synovium, harmony was used to integrate across samples and the study of origin^56^. Leiden clustering was applied to derive broad cell populations including: B cells, plasma cells, T cells, myeloid cells, and stromal cells.

These broad cell populations were then further clustered as described above, with tailored PCAs and n_neighbors as per dataset complexity in addition to harmony for batch correction. In instances where individual cell clusters in these partitioned datasets demonstrated biological anomalies (such as high B cell marker gene expression in a distinct T cell cluster), the Scrublet score was used to assess the likelihood that this cluster could be a doublet, in which case these cells would be removed from the analysis. The Wilcoxon rank-sum test was used to conduct differential expression between clusters to derive marker genes. False discovery rate (FDR)-adjusted *P*-value < 0.05 considered significant for marker genes and all other analyses unless otherwise specified.

### Derivation of the inflammation score

The inflammation score is a composite gene score. We identified genes differentially expressed between histologically inflamed (as per Nancy index) IBD resections to non-inflamed/non-IBD gut tissue following multiple comparison correction using DESeq2^Ref^^27,57^ **(Supplementary Table 4**). Data derived from TAURUS were pseudobulked (sum) at the sample level. We then used the aforementioned list of differentially expressed genes as a gene signature and applied the *enrichIt* function from the *escape* package^58^. The score was then scaled between 0-10. This resulted in a vector representing enrichment of the inflammation score on a per sample basis. The highest inflammation score in the healthy sample was selected as a heuristic cut off for the inflammation score. This corresponds to the 90^th^ percentile of inflammation score found in macroscopically non-inflamed samples.

### Treatment response criteria

Patients with CD who experienced 30% decrease in HBI and changed to no macroscopic ulceration following treatment were considered to be responders. Patients with UC who experienced 30% decrease in SSCAI, Nancy index and UCEIS were considered to be responders. If these outcome measures were incongruent, the inflammation score in the post treatment samples were examined. Patients in whom any post-treatment samples remained inflamed as per the inflammation score (>6.5) were categorised as non-responders. If all post-treatment samples were below the inflammation score threshold (<6.5), patients would be considered responders. Any patient who required surgical intervention, or necessitated change in therapy due to disease activity was categorised as a non-responder. For RA, we used a EULAR good or moderate response to define binary response in RA^59^.

### Differential abundance analysis

Differential abundance was carried out using MASC with nested random effects accounting for multiple samples per patient, and covariates including treatment status^60,61^. Differential abundance was conducted in two ways:

i. To detect cell state specific changes in inflammation, comparison across CD and UC, treatment response associations at baseline and effect of treatment, cell state abundance was analysed as proportion of the ‘low’ resolution category **(Extended Data Fig. 1b)**
ii. To detect compartment-specific changes following treatment across responders and non-responders, compartment abundance was analysed as proportion of the entire sample.

### PROGENy analysis

To quantify TNF signalling, we applied PROGENy to our dataset^38^. Each cell received a score for each of 14 pathways including TNF signalling. Linear mixed effects model using the *lmer* function as part of the *lmerTest* package was used to test for association between TNF signalling scores before and after anti-TNF therapy with the patient variable accounted as random effects.

### scRNA-seq differential expression and pathway analysis

Pseudobulked profiles were generated at the compartment level for differential expression analysis. Comparisons between ileum and colon were performed using *limma-voom* with *duplicateCorrelation* to account for multiple samples per patient^62^. Linear model was fit using *lmFit*, and moderated t-statistics as well as associated *P*-values were generated using the *ebayes* function.

To longitudinally monitor compartmental changes following therapy, we applied *glmmSeq* to paired samples^42^. Counts Per Million (CPM) normalisation was applied to pseudobulked profiles. Given that *glmmSeq* used negative binomial models, we generated estimates of the common, trended and genewise dispersions across all genes using *estimateDisp* function in *edgeR*^63^. An interaction term of treatment (pre/post) by response (responder/non-responder) was used alongside a nested random effects design to account for multiple samples from the same patient. For overrepresentation analysis, *enrichGO* from *clusterProfiler* was used^64^. All genes tested were used as the background genes i.e. the gene universe. Genes with positive fold change in non-responders as well as q-value < 0.05 for treatment:response were tested for Gene Ontology (GO) term enrichment.

### Identification of gene expression programmes (GEPs) by consensus non-negative matrix factorisation (cNMF)

We leveraged cNMF to complement the Leiden-based clustering approach to simultaneously capture functional programmes, and activation states in addition to cell identity^40^. cNMF was iteratively applied to broad categories of cell types as identified with Leiden clustering. In the gut, this consisted of: B cells, plasma cells, CD4^+^ T cells, CD8^+^ T cells, myeloid cells (monocytes, macrophages and dendritic cells, mast cells), stromal cells (fibroblasts and pericytes), myofibroblasts, endothelial cells, colonic epithelial cells, ileal epithelial cells, glial cells and innate lymphoid cells. In the synovium, this consisted of: B cells, plasma cells, T cells, myeloid, and the stroma.

Briefly, we applied cNMF to a count matrix, *N* (cells) x *M* (genes) to derive two matrices: *k* (GEP) x *M* (genes), and, *N* (cells) x *k* (GEP) with the usage of each GEP for each cell^40^. Selection of *k* was dependant on several factors including prioritising solutions that were biologically meaningful according to top weighted genes, factorisation stability as determined by the silhouette score and minimisation of the Frobenius reconstruction error. Consensus solutions were then filtered for outliers through inspections of distances between components and their nearest neighbours through a histogram. Genes statistically associated with each GEP was identified using multiple least squares regression of normalised (z-scored) gene expression against the consensus GEP usage matrix. Overrepresentation analysis for all GEPs were conducted through using GOATOOLS with top 150 weighted genes^65^ as input and all genes in the relevant matrix as the gene universe.

### Identification of hubs and calculating NMF transcriptional programme activity

Hubs were identified through analysis of covarying GEPs in inflamed samples for CD and UC separately^66^. Programme activity was calculated for every GEP according to the cell type in which the GEP was initially discerned. For the gut, this was restricted to the main non-epithelial cell types; B cells, plasma cells, CD4^+^ T cells, CD8^+^ T cells, myeloid cells (monocytes, macrophages and DC), and stromal cells (fibroblasts and pericytes). For the synovium, this included B cells, plasma cells, T cells, myeloid, and the stroma.

As previously demonstrated, we summarised programme activity for each GEP across individual samples^66^. We calculated GEP expression across five quantiles (0.25, 0.5, 0.75, 0.95, 0.99) individually for each sample. For each quantile, a Pearson correlation co-efficient (*R*) was derived for each pair of GEPs across all samples. The correlation was Fisher transformed and the mean of these correlations were used as a test statistic. We compared *R* against a null distribution derived through permuting the sample identity 10,000 times keeping cell type constant. A *P*-value was generated through counting how often the permuted *R* value was above and below the true *R* value. The minimum count was scaled by two and designated the *P*-value statistic. Multiple comparisons were corrected at Benjamini-Hochberg FDR of 10%. We derived an adjusted *R* value by calculating the difference of mean true *R* values and the mean of permuted *R* values.

Significant Fisher transformed associations, *R* (edges) and their constituent GEPs (nodes) were used to create a signed weighted network. Hubs within this network were detected using a module detection algorithm used for signed graphs^67^. This was applied by resolution parameter in the range of 0.001 to 0.2, and tau=0.2. This method was iteratively applied, and hubs split if they were larger than three nodes, and improved modularity of the solution.

### Testing GEP enrichment in inflammation

In order to capture GEPs that are active even in a small number of cells, we calculated the mean of programme activity values at five percentiles (0.25 0.5, 0.75, 0.95, 0.99). Linear mixed effects model using the *lmer* function as part of the *lmerTest* package was used to test for enrichment of GEPs in inflammation in the gut. Association between mean GEP expression and inflammation status was tested with covariates including patient and treatment. Hubs in IBD were deemed to be inflammatory if more than half of the constituent GEPs in a particular hub was enriched in inflammation. Hubs in RA were deemed to be inflammatory if more than half of the constituent GEPs in a particular hub was positively correlated with the proportion of CD45^+^ cells in samples^45^.

### Projection of GEPs to bulk RNA sequencing/microarray data

As outlined above, cNMF yields a *k* (GEP) x *M* (genes) matrix, henceforth referred to as *H.* The gene expression matrix from the relevant microarray/bulk RNA sequencing data were subsetted to genes shared with *H*. NMF was initialised with *H* and the gene expression matrix to generate the projected component matrix, *W* (samples x *k*). The NMF implementation used was *sklearn.decomposition.non_negative_factorization*.

### Processing bulk RNA sequencing data from R4RA

FASTQ files generated from the R4RA trial were downloaded from EMBL-EBI (E-MTAB-11611). FASTQ files were trimmed to remove low-quality reads using trimgalore (0.6.6) in paired mode. FASTQ files were aligned to the human genome (GRCh38, Ensembl release 101) using STAR (2.7.3a). Gene counts were summarised using featureCounts (Subread v2.0.1). Raw counts were RPKM-normalised using edgeR functions *calcNormFactors* (TMM) and *rpkm*.

### Multiplexed imaging using Cell DIVE

#### Slide clearing and blocking

4 μM Formalin-Fixed Paraffin Embedded (FFPE) biopsy tissues slides (CD or UC) were deparaffinised and rehydrated. The slides were then permeabilised for 10 min in 0.3% Triton X-100 and washed further in 1X PBS. Antigen retrieval was performed using the NxGen decloaking chamber (Biocare Medical, Pacheco, CA, USA) in boiling pH6 Citrate (Agilent, S1699) and pH9 Tris-based antigen retrieval solutions for 20 min each. Tissue slides were blocked in 1X PBS with a 3% BSA (Merck, A7906), 10% Donkey serum (Bio-Rad, C06SB) solution for 1 h at room temperature. Slides were washed in 1X PBS for 10 min and then stained with DAPI (Thermo, D3571) for 15 min. Slides were washed in 1X PBS for 5 min and coverslipped with mounting media (50% glycerol – Sigma, G5516 and 4% propyl gallate – Sigma, 2370).

#### Scan plan and background acquisition

The GE Cell DIVE system was used to image all FFPE slides. A scan plan was acquired at 10X magnification to select regions of interest followed by imaging at 20X to acquire background autofluorescence and generate virtual H&E images. Background imaging is used to subtract autofluorescence from all subsequent rounds of staining. Slides were decoverslipped in 1X PBS prior to staining.

#### Staining and bleaching

Multiplexed imaging consisted of staining for the following protein markers: CD68, CD3, CCL19, CD8A, CK8, CD4, CXCL13, CD20, CD208, CXCL9, S100A9, KI67, MPO, GZMB, CD66B, CD14, MZB1, CK8, COL1A1, CCR7, CD11C, CD40, PD1. Each staining round consisted of a mix of three antibodies prepared in blocking buffer (PBS, 3% BSA, 10% donkey serum). The initial round used primary antibodies which were incubated overnight at 4°C followed by washes in 1X PBS and 0.05% Tween20 (Sigma, P9416). Secondary antibodies raised in Donkey were then incubated for an additional hour at room temperature which were either conjugated to Alexa Fluorophore 488, 555 or 647 (Invitrogen). Each subsequent staining round used directly conjugated antibodies to either of these dyes and were incubated overnight at 4°C. Antibodies manually conjugated were purchased in a BSA-AZIDE free format and conjugated using antibody labelling kit (Invitrogen). Fluorophores were bleached between each staining round using NaHCO_3_ (0.1 M, pH 11.2; Sigma, S6297) and 3% H_2_O_2_ (Merck, 216763). Fresh bleaching solutions were prepared and slides were bleached two times (15 min each) with a 1 min 1X PBS wash in between bleaching rounds. Slides were re-stained for DAPI for 2 min and washed in 1X PBS for 5 min before imaging the dye-inactivated round as the new background round (for subsequent background subtraction). DAPI staining between imaging rounds assists in image registration and alignment. Slides were multiplexed with the next panel of three markers with iterative staining, bleaching and imaging.

## Acknowledgements

TT and CDB are supported by the Kennedy Trust for Rheumatology Research through the Arthritis Therapy Acceleration Programme (ATAP). TT was supported by Celsius Therapeutics. Computational aspects of this research were supported by the Wellcome Trust Core Award Grant Number 203141/Z/16/Z with additional support from the NIHR Oxford BRC. The views expressed are those of the author(s) and not necessarily those of the NHS, the NIHR or the Department of Health. CRG and CDB are supported by the NIHR Oxford Biomedical Research Centre (BRCRCF10-04) and Cartography Consortium funding from Janssen Biotech, Inc. MP is supported by the UK Medical Research Council (MR/W025981/1). MF is supported by the Wellcome Trust (225928/Z/22/Z), Crohn’s and Colitis UK M2021/2, and the Medical Research Council (MR/W025981/1). MF was supported by Oxford-UCB and Oxford-Janssen fellowships. AV is supported by a Kennedy Trust Prize Studentship. CAD is supported by a Wellcome Trust and Royal Society (204290/Z/16/Z), the UK Medical Research Council (MR/T030410/1), the Rosetrees Trust (R35579/AA002/M85-F2), Cartography Consortium funding from Janssen Biotech, Inc and the NIHR Oxford Biomedical Research Centre, Inflammation Across Tissues and Cell and Gene Therapy Themes. Celsius Therapeutics funded the generation of the single-cell transcriptomics data. Ginevra Botta provided valuable support for the execution of the study. Sara Ruzycky, Jacki Growe and Cindy Hession generated transcriptomics data. Colin Smith, Gerard Honig and Malcolm Daniels supported the collaboration between Celsius Therapeutics and Oxford University. Schematics in figures have been produced with the aid of BioRender.com

## Author contributions

Conceptualization was the responsibility of TT, RP, RK, NSpies, ST, HHU, CAD, CDB. Methodology was undertaken by all authors. Computational analysis was undertaken by TT, CRG, MF, AJ, AV, NS, NSpies, CAD. Validation was performed by TT, MP, MF, DA, CAD. Resources were the responsibility of TT, ST, HHU, CAD, CDB. Data curation was performed by TT, CRG, NSpies, CAD. Writing of the original draft was done by TT, HHU, ST, CAD, CDB. Writing, review and editing were done by all authors listed. ST, HHU, CAD, CDB supervised the project.

## Data availability

All raw and processed data will be available upon acceptance via Human Cell Atlas, EGA and Zenodo. A cell browser website will be available to visualise our data and results.

## Code availability

All source code will be available on GitHub upon acceptance of this paper. Supplementary information is available for this paper.

## Figure legends

**Extended Data Fig. 1.**
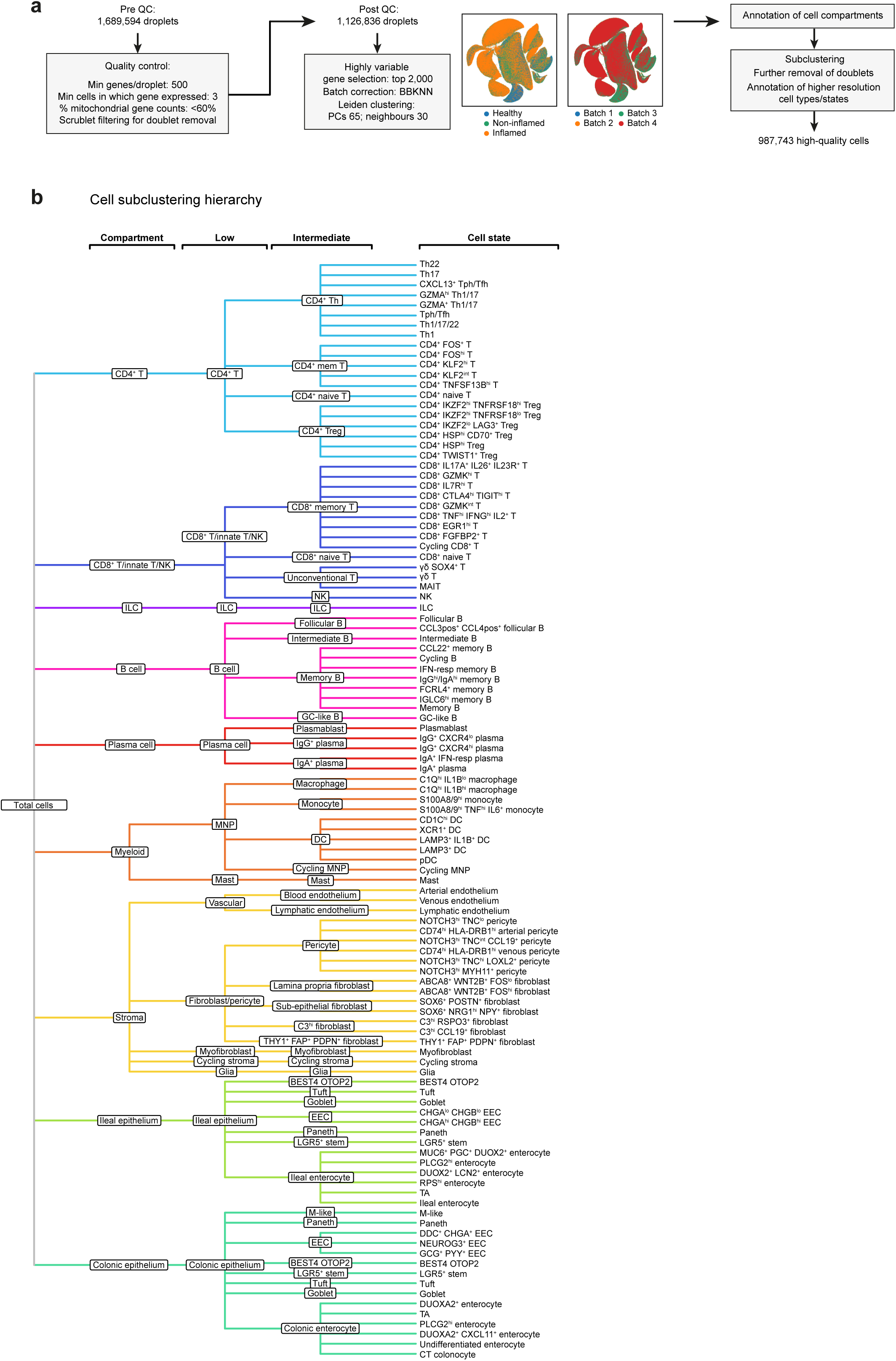
Sample processing and annotation hierarchy. **a,** Schematic showing bioinformatic pre-processing strategy for gut samples. Panpipes pipeline was used for pre-processing^53^. Uniform manifold approximation and projection (UMAP) visualisations show the cellular landscape of gut samples coloured by inflammation status, and batch. See **Methods** for more details. **b,** Hierarchy shows annotation across increasing cell type resolution: compartment, low, intermediate and cell state. Colono, colonocyte; DC, dendritic cell; EEC, enteroendocrine cell; entero, enterocyte; fibro, fibroblast; GC, germinal centre; hi, high; IFN-resp, interferon-responsive; ILC, innate lymphoid cell; lo, low; macro, macrophage; MAIT, mucosal-associated invariant T; MNP, mononuclear phagocyte; mono, monocyte; NK, natural killer; PC: principal components; pDC, plasmacytoid dendritic cell; peri, pericyte; TA, transit-amplifying; Tfh, CD4^+^ follicular helper T cell; Th, CD4^+^ helper T cell; Tph, CD4^+^ peripheral helper T cell; Treg, regulatory T cell; Undiff, undifferentiated.

**Extended Data Fig. 2.**
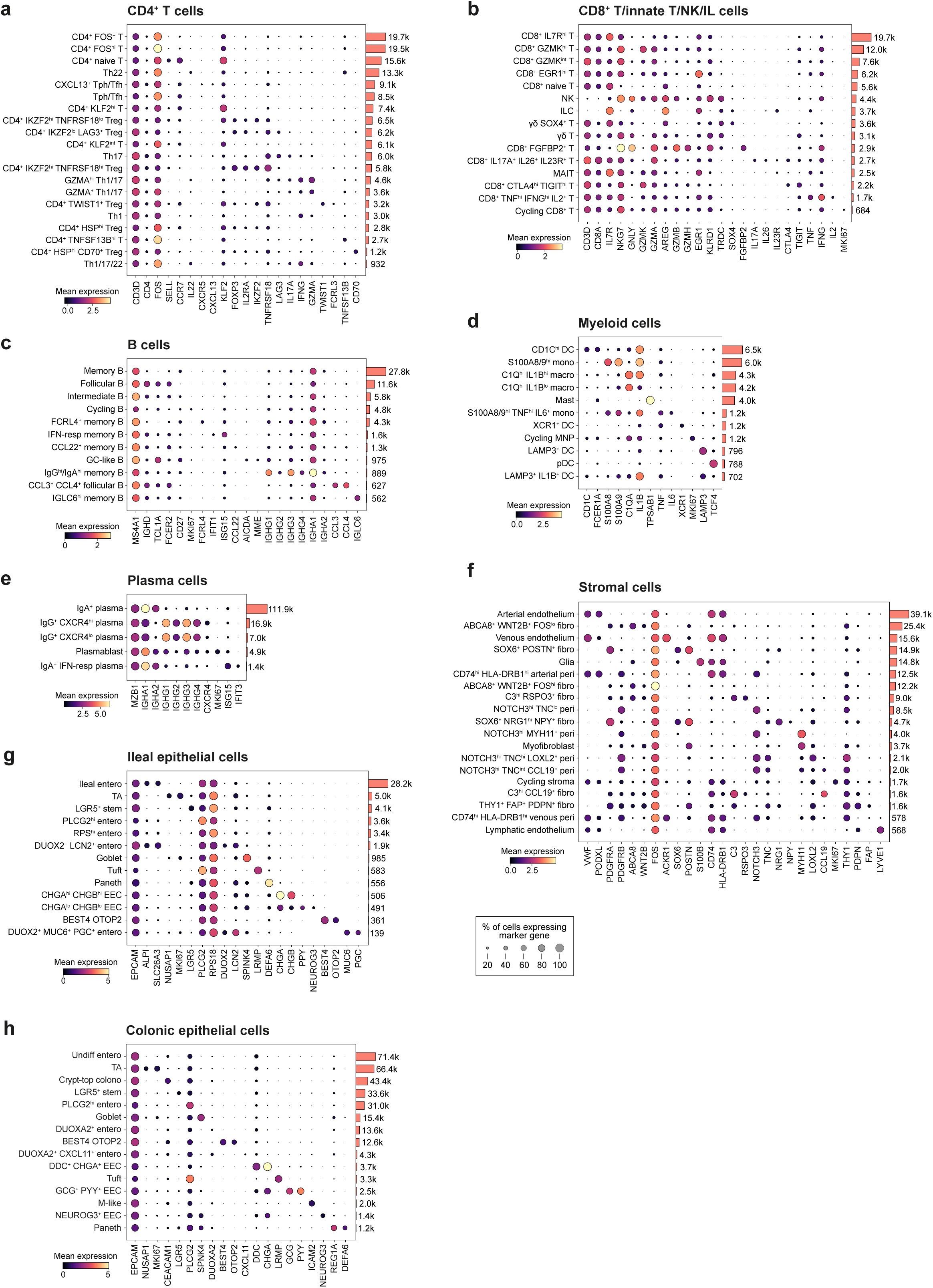
Marker genes of cell states in the gut. Dotplot showing expression of marker genes of cell states in the scRNA-seq dataset: **(a)** CD4^+^ T cell, **(b)** CD8^+^ T/innate T/NK/IL cell, **(c)** B cell, **(d)** myeloid cell, **(e)** plasma cell, **(f)** stromal cell, **(g)** ileal epithelial cell and **(h)** colonic epithelial cell. Genes relate to **Supplementary Table 2**. Colono, colonocyte; DC, dendritic cell; EEC, enteroendocrine cell; entero, enterocyte; fibro, fibroblast; GC, germinal centre; hi, high; IFN-resp, interferon-responsive; ILC, innate lymphoid cell; lo, low; macro, macrophage; MAIT, mucosal-associated invariant T; MNP, mononuclear phagocyte; mono, monocyte; NK, natural killer; pDC, plasmacytoid dendritic cell; peri, pericyte; TA, transit-amplifying; Tfh, CD4^+^ follicular helper T cell; Th, CD4^+^ helper T cell; Tph, CD4^+^ peripheral helper T cell; Treg, regulatory T cell; Undiff, undifferentiated.

**Extended Data Fig. 3.**
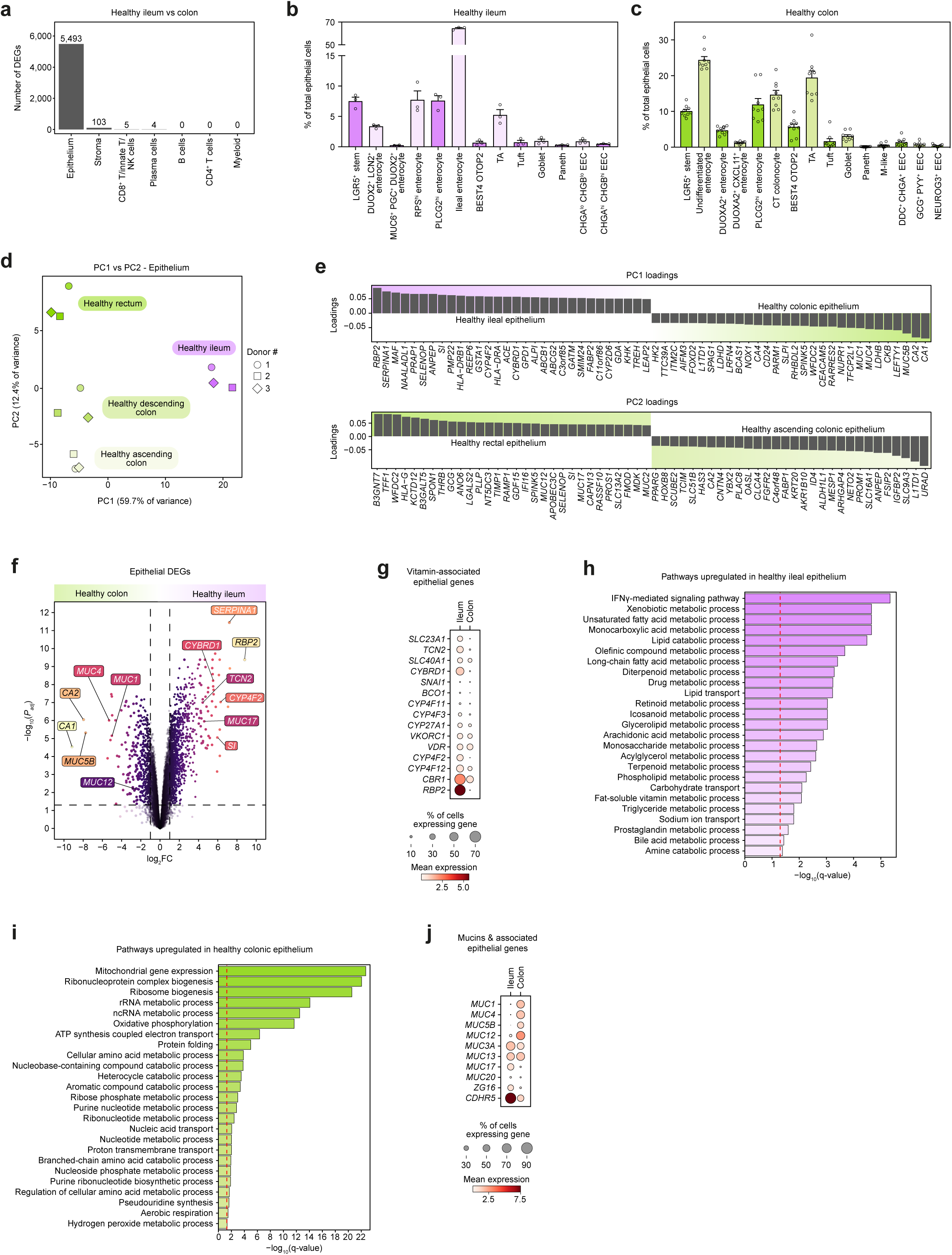
Differences between the healthy ileum and colon. **a,** Barplot summarising number of differentially expressed genes (DEGs) (*P_adj_*< 0.05) comparing healthy ileum (three samples) to healthy colon (nine samples) in three patients in each cell compartment. Limma-voom with *DuplicateCorrelation* used to adjust for multiple samples per patient^62^. **b,c,** Cell state distribution within the epithelial compartment in **(b)** ileum and **(c)** colon displayed on a bar plot. Error bar indicates standard error of mean. **d,** *prcomp* from base *R* used to conduct PCA on CPM normalised and log-transformed read counts. Samples in context of principal components (PC) 1 and 2 along with associated percentage of variation explained. **e,** Loadings of genes associated with PC1 and PC2 shown in the bar plots. **f,** Volcano plot showing results of differential expression between ileum and colon in the epithelial compartment. Dashed lines demarcate *P_adj_* =0.05 and log_2_ fold change (FC)=0.5. **g,** Relative expression of vitamin-associated epithelial genes differentially expressed between ileum and colon shown in dotplot. Full results can be found in **Supplementary Table 3**. **h,i,** Over-representation analysis was performed by using the enrichGO function from *clusterProfiler*^64^. All genes significantly associated with **(h)** ileum and **(i)** colon respectively tested for overrepresentation using gene ontology (GO) biological process gene sets. Red dashed line indicative of q-value=0.05. **j,** Relative expression of mucin and mucin-associated genes differentially expressed between ileum and colon shown in dotplot. Full results can be found in **Supplementary Table 3**.

**Extended Data Fig. 4.**
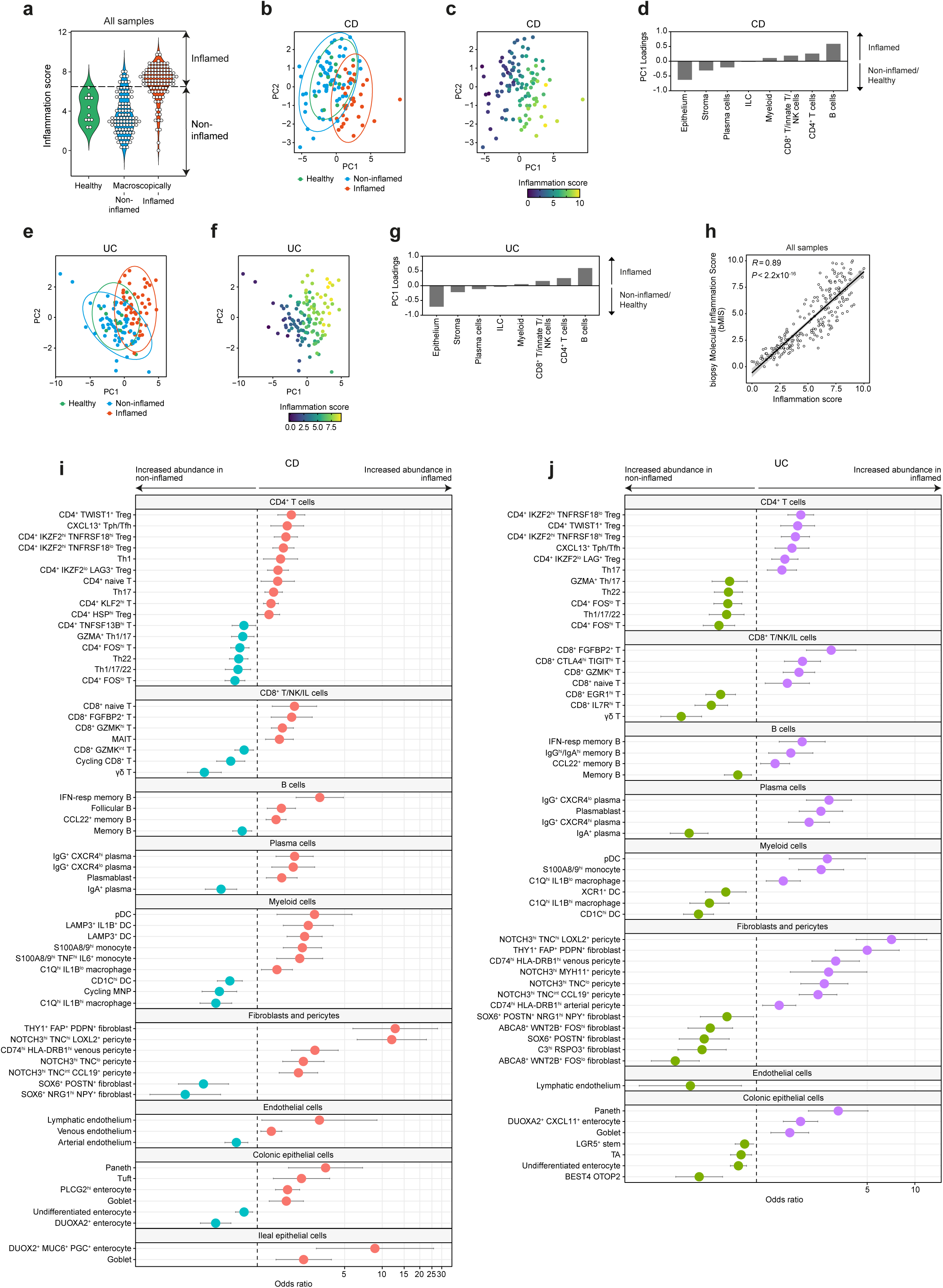
The inflammation score in context of CD and UC. **a,** Violin plot showing the distribution of the inflammation score across healthy and macroscopically non-inflamed, as well as inflamed samples. **b,** PCA examining compartment abundance as a proportion of sample in CD. **c,** PCA of samples with CD with inflammation score plotted as a quantitative variable. **d,** PC1 loadings associated with cell compartment in samples with CD. **e,** PCA examining compartment abundance as a proportion of sample in UC. **f,** PCA of samples with UC with inflammation score plotted as a quantitative variable **g,** PC1 loadings associated with cell compartment in samples with UC **h,** Spearman correlation between inflammation score per sample and the recently described biopsy molecular inflammation score (bMIS)^29^ **i, j,** Differential abundance of cell states in CD **(i**) and UC **(j**) comparing non-inflamed to inflamed tissue. Error bars show 95% confidence interval. DC, dendritic cell; hi, high; IFN-resp, interferon-responsive; ILC, innate lymphoid cell; lo, low; macro, macrophage; MAIT, mucosal-associated invariant T; MNP, mononuclear phagocyte; mono, monocyte; NK, natural killer; pDC, plasmacytoid dendritic cell; TA, transit-amplifying; Tfh, CD4^+^ follicular helper T cell; Th, CD4^+^ helper T cell; Tph, CD4^+^ peripheral helper T cell; Treg, regulatory T cell.

**Extended Data Fig. 5.**
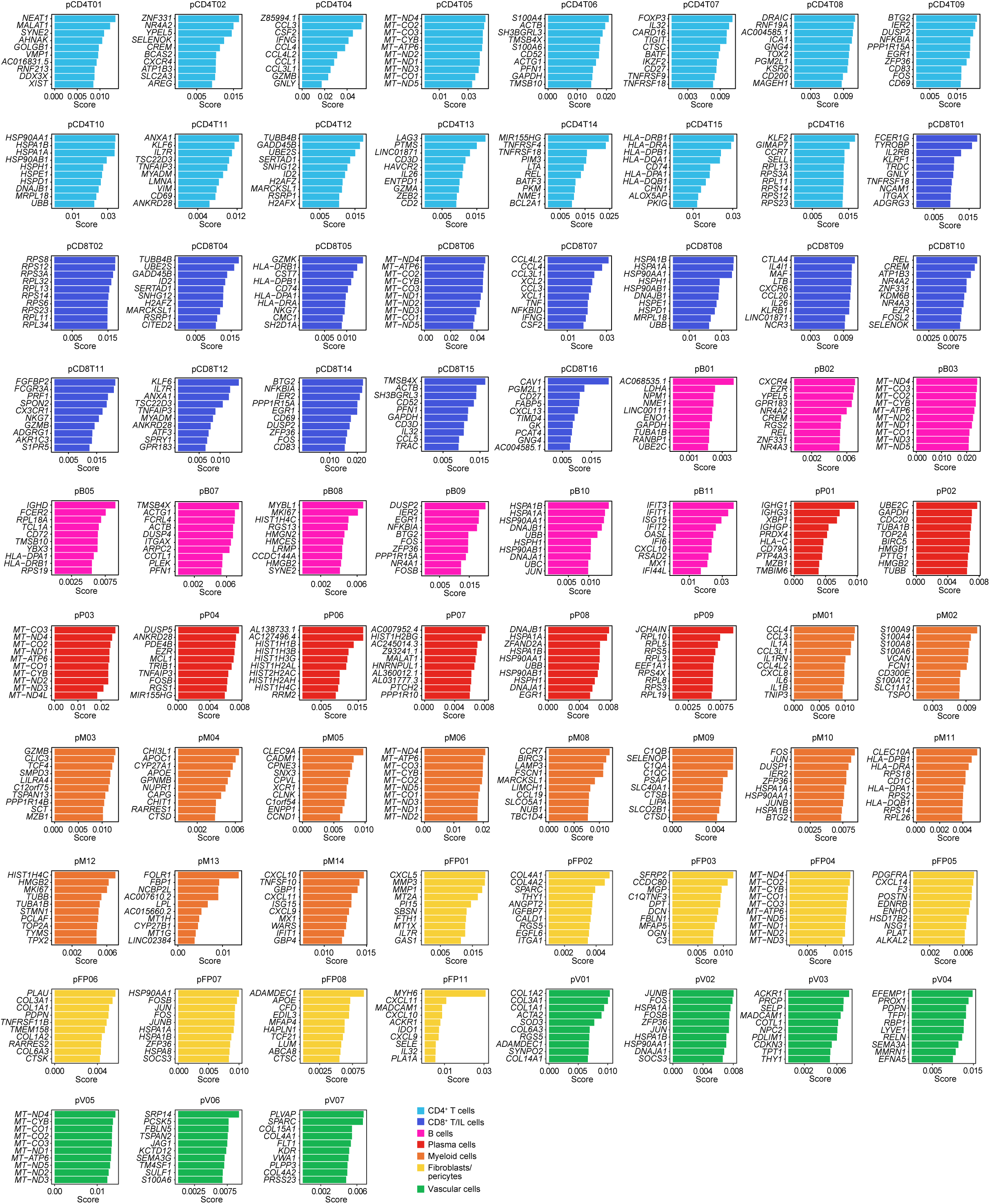
Top weighted genes of GEPs in the gut. Weighted genes for each gene expression programme (GEP) derived from the gut. cNMF was run separately in: CD4^+^ T, CD8^+^ T, B, plasma cells, myeloid cells (monocytes, macrophages and DC), vascular cells, and fibroblasts and pericytes. See **Supplementary Table 5** for full list of weighted genes, as well as results of overrepresentation analysis. See **Supplementary Table 6** for results of enrichment testing of GEPs in inflammation. pB: B cell GEP; pCD4T: CD4^+^ T cell GEP; pCD8T: CD8^+^ T cell/NK GEP; pFP: fibroblast and pericyte GEP; pM: myeloid cell GEP; pP: plasma cell GEP; pV: vascular cell GEP.

**Extended Data Fig. 6.**
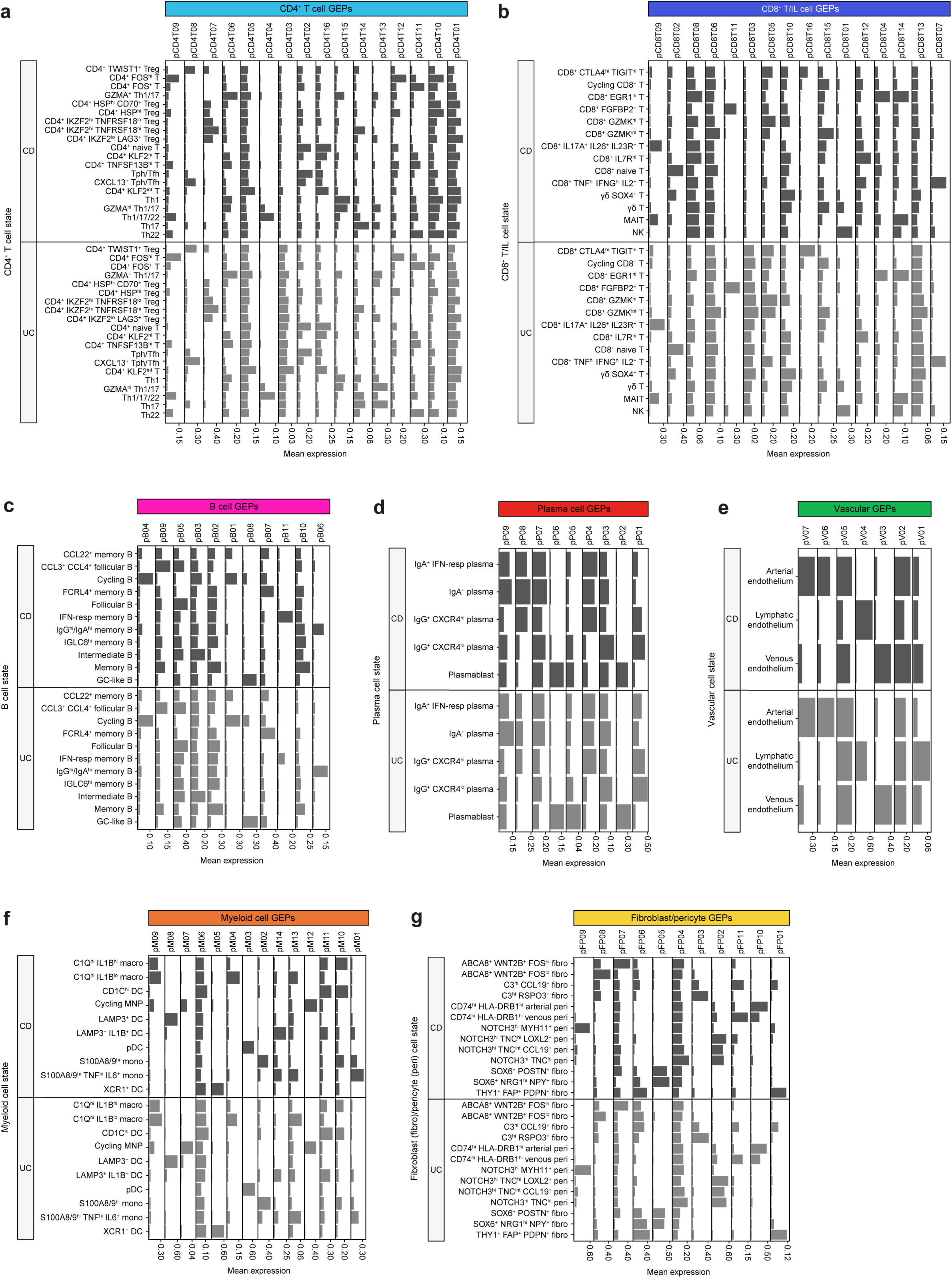
Enrichment of GEPs across cell states in the gut. cNMF was used to derive GEP score for individual cells from inflamed samples with CD and UC in **(a)** CD4^+^ T, **(b)** CD8^+^ T, **(c)** B, **(d)** plasma, **(e)** vascular, **(f)** myeloid, and **(g)** fibroblast and pericyte cells. Mean expression of GEP quantified per cell state is plotted. pB: B cell GEP; pCD4T: CD4^+^ T cell GEP; pCD8T: CD8^+^ T cell/NK GEP; pFP: fibroblast and pericyte GEP; pM: myeloid cell GEP; pP: plasma cell GEP; pV: vascular cell GEP. DC, dendritic cell; fibro, fibroblast; GC, germinal centre; hi, high; IFN-resp, interferon-responsive; ILC, innate lymphoid cell; lo, low; macro, macrophage; MAIT, mucosal-associated invariant T; MNP, mononuclear phagocyte; mono, monocyte; NK, natural killer cell; pDC, plasmacytoid dendritic cell; peri, pericyte; Tfh, CD4^+^ follicular helper T cell; Tph, CD4^+^ peripheral helper T cell; Th, CD4^+^ T helper cell; Treg, CD4^+^ regulatory T cell.

**Extended Data Fig. 7.**
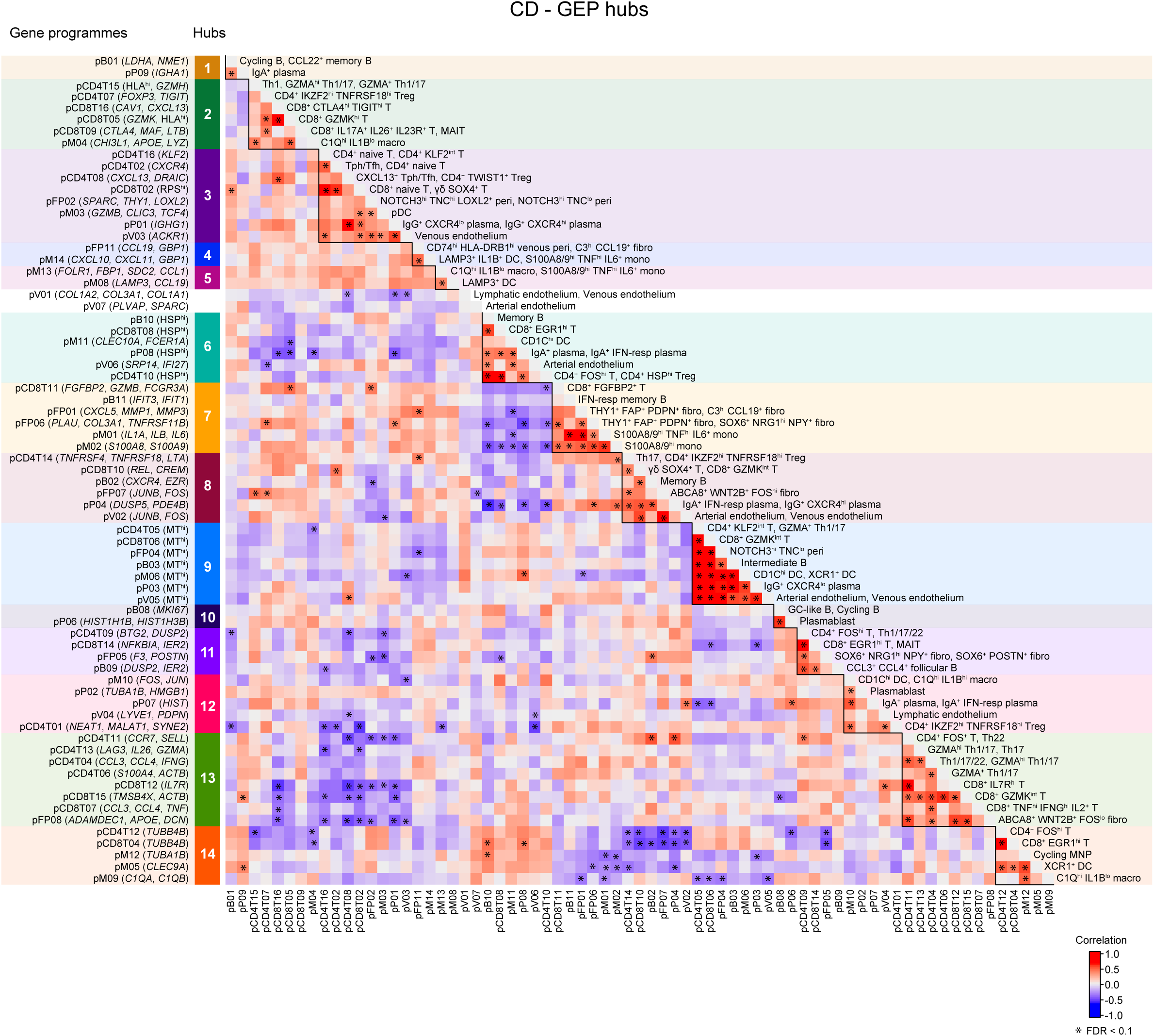
Covarying GEPs in inflamed samples with CD. Correlogram demonstrating significant correlations (asterisks: FDR< 0.1) between GEPs across cell compartments in inflamed samples with CD. Lines demarcate hubs. A module detection algorithm used for signed graphs was leveraged to detect hubs from a graph consisting of significantly correlated GEPs (nodes) and associated fisher-transformed correlations (edges)^67^. DC, dendritic cell; fibro, fibroblast; GC, germinal centre; hi, high; HSP, heat-shock proteins; IFN-resp, interferon-responsive; ILC, innate lymphoid cell; lo, low; macro, macrophage; MAIT, mucosal-associated invariant T; MNP, mononuclear phagocyte; mono, monocyte; MT^hi^, mitochondrial-high; NK, natural killer cell; pB, B cell GEP; pCD4T, CD4^+^ T cell GEP; pCD8T, CD8^+^ T cell/NK GEP; pDC, plasmacytoid dendritic cell; peri, pericyte; pFP, fibroblast and pericyte GEP; pM, myeloid cell GEP; pP, plasma cell GEP; pV, vascular cell GEP; RPS^hi^, ribosomal protein S-high; Tfh, CD4^+^ follicular helper T cell; Tph, CD4^+^ peripheral helper T cell; Th, CD4^+^ T helper cell; Treg, CD4^+^ regulatory T cell.

**Extended Data Fig. 8.**
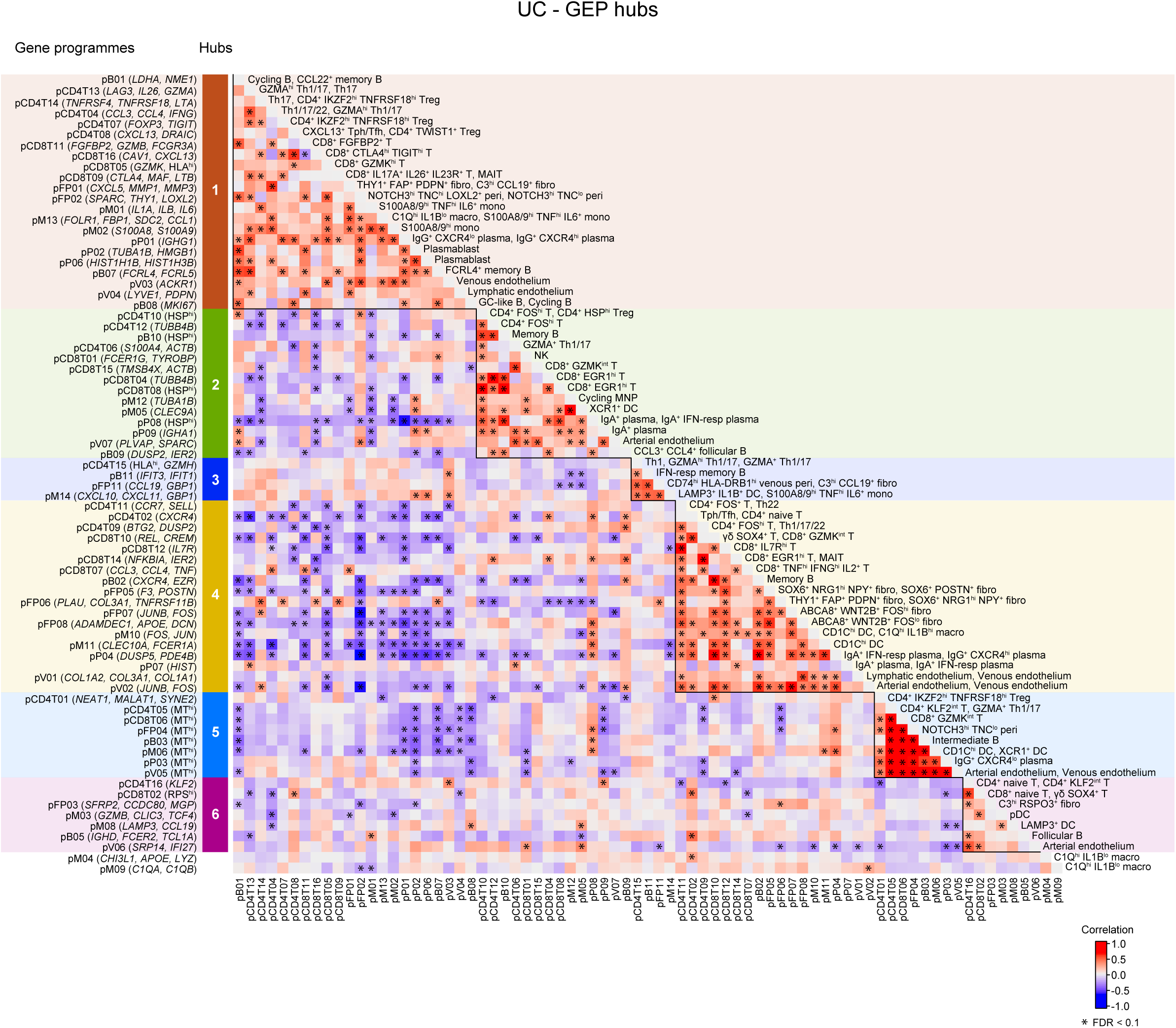
Covarying GEPs in inflamed samples with UC. Correlogram demonstrating significant correlations (asterisks: FDR< 0.1) between GEPs across cell compartments in inflamed samples with UC. Lines demarcate hubs. A module detection algorithm used for signed graphs was leveraged to detect hubs from a graph consisting of significantly correlated GEPs (nodes) and associated fisher-transformed correlations (edges)^67^. DC, dendritic cell; fibro, fibroblast; GC, germinal centre; hi, high; HSP, heat-shock proteins; IFN-resp, interferon-responsive; ILC, innate lymphoid cell; lo, low; macro, macrophage; MAIT, mucosal-associated invariant T; MNP, mononuclear phagocyte; mono, monocyte; MT^hi^, mitochondrial-high; NK, natural killer cell; pB, B cell GEP; pCD4T, CD4^+^ T cell GEP; pCD8T, CD8^+^ T cell/NK GEP; pDC, plasmacytoid dendritic cell; peri, pericyte; pFP, fibroblast and pericyte GEP; pM, myeloid cell GEP; pP, plasma cell GEP; pV, vascular cell GEP; RPS^hi^, ribosomal protein S-high; Tfh, CD4^+^ follicular helper T cell; Tph, CD4^+^ peripheral helper T cell; Th, CD4^+^ T helper cell; Treg, CD4^+^ regulatory T cell.

**Extended Data Fig. 9.**
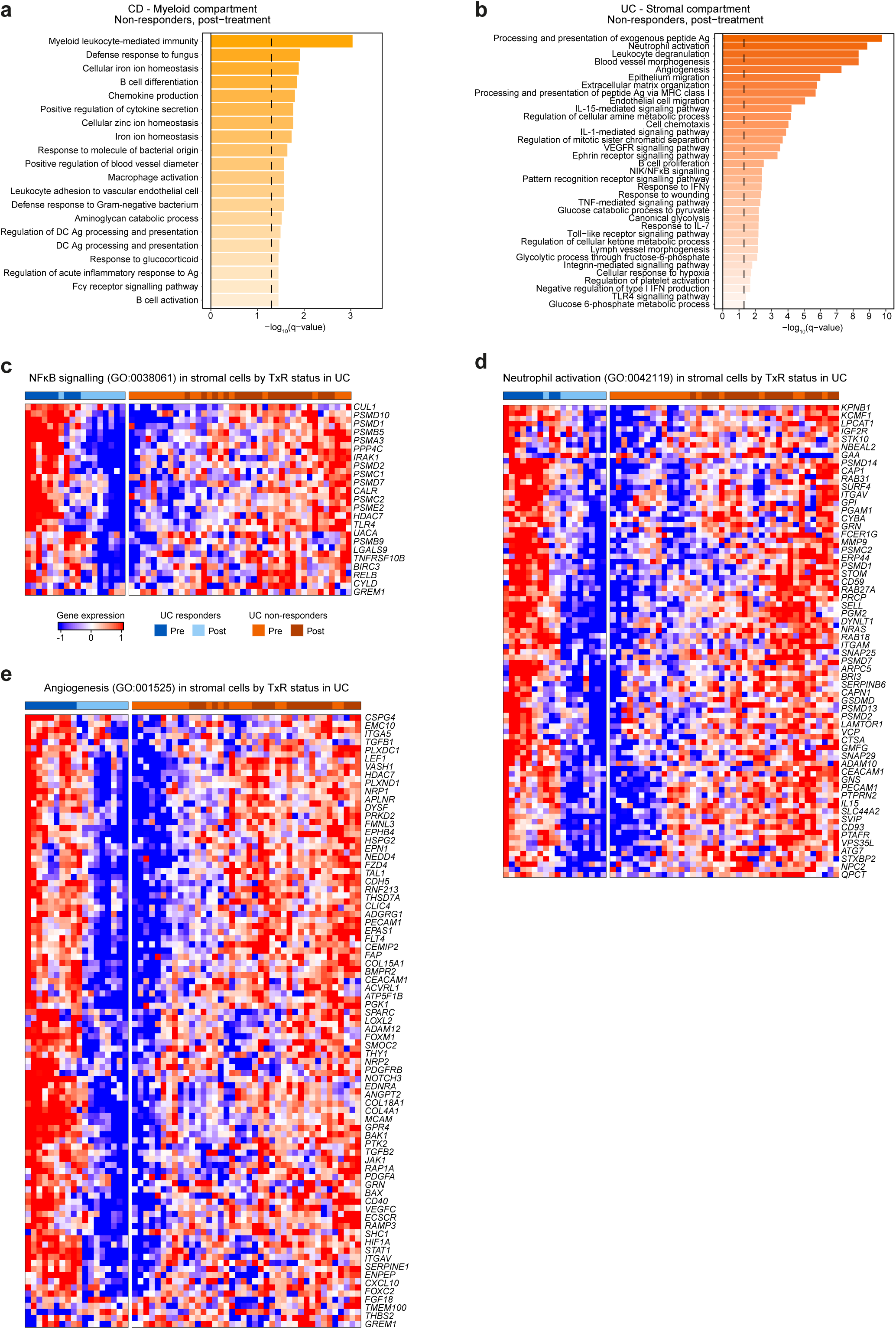
GO over-representation analysis of changes following anti-TNF therapy in the myeloid and stromal compartments of non-responding patients with CD and UC. **a,b,** Over-representation analysis was performed by using the *enrichGO* function from *clusterProfiler*^64^. Genes with positive fold change in non-responders with significance (*P_adj_* <0.05) for treatment:response (TxR) were tested for GO term enrichment and shown in barchart for **(a)** myeloid compartment in CD **(b)** stromal compartment in UC. Dashed line indicates q-value = 0.05. **c-e)** Following over-representation analysis in the stromal compartment of non-responders with UC, genes differentially expressed and associated with specific GO terms, **(c)** GO:0038061 **(d)** GO:0042119 **(e)** GO: 0001525 were examined in the paired samples included in the differential expression analysis. Genes were log transformed, and scaled from raw counts. The heatmap was then generated using *ComplexHeatmap* and split into responders and non-responders with column-wise and row-wise hierarchical clustering. Expression was scaled between −1 to 1.

**Extended Data Fig 10.**
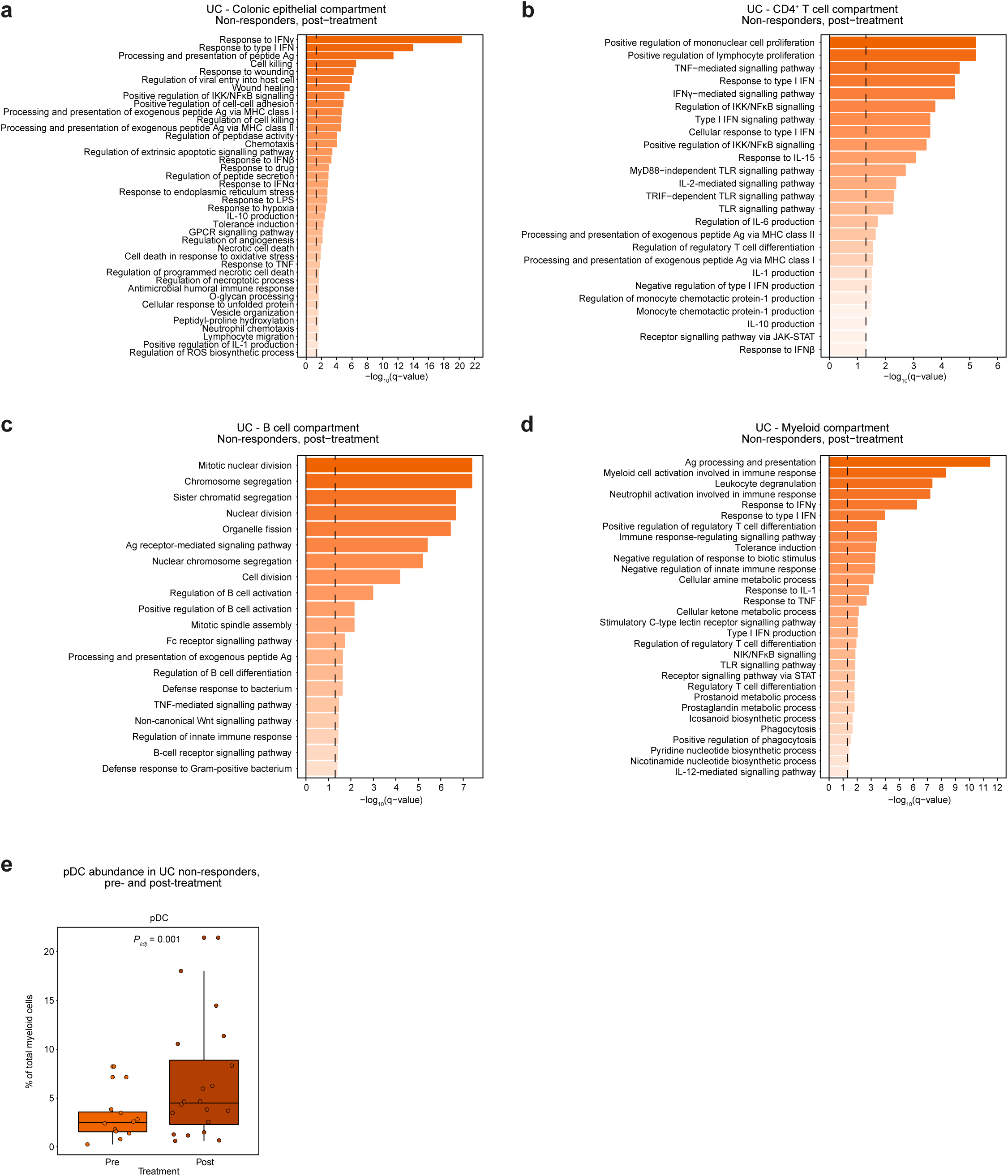
Changes in colonic epithelium, CD4^+^ T cells, B cells and myeloid cells following treatment with anti-TNF in non-responding patients with UC. **a-d,** Over-representation analysis was performed by using the *enrichGO* function from *clusterProfiler*^64^. Genes with positive fold change in non-responders with significance (*P_adj_* < 0.05) for treatment:response were tested for GO term enrichment and shown in bar chart for **(a)** colonic epithelium **(b)** CD4^+^ T cells **(c)** B cells and **(d)** myeloid cells in UC. **e,** Boxplot showing proportion of plasmacytoid DC (pDC) cell state out of the proportion of mononuclear phagocytes before and after treatment with anti-TNF in non-responders with UC. Only paired samples included in the analysis. Differential abundance tested using MASC with nested random effects accounting for multiple samples per patient (*P_adj_* < 0.05).

**Extended Data Fig. 11.**
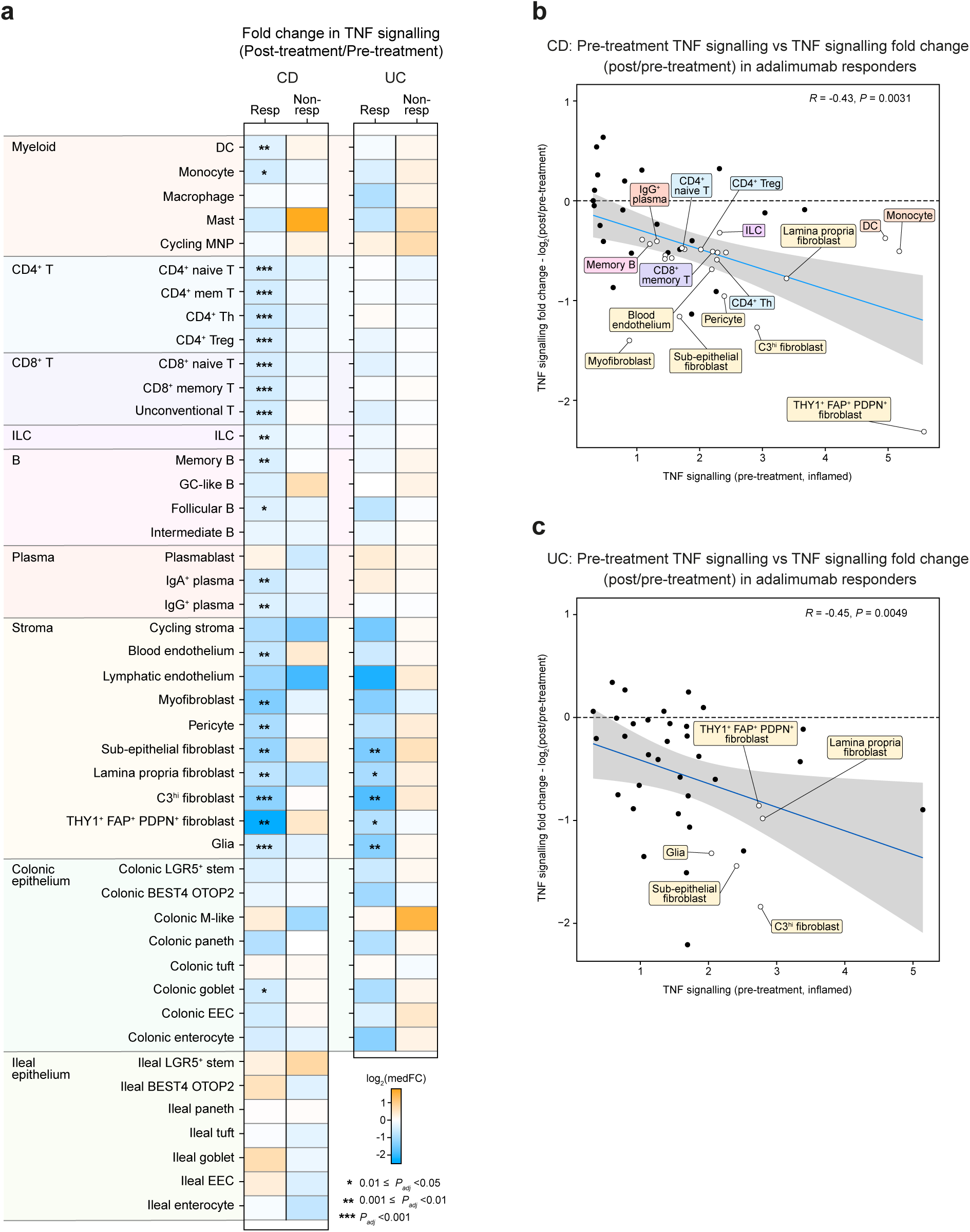
Characterisation of the TNF pathway following anti-TNF therapy in IBD. **a,** PROGENy was used to calculate TNF signalling scores at a per cell level^38^. The 75^th^ percentile score for TNF signalling in each of cell types at the ‘intermediate’ level of resolution was taken to be representative of individual samples. Only paired samples were used to calculate median fold change (medFC) in responders (Resp) and non-responders (Non-resp) with significance testing using *lmer* function as part of the *lmerTest* package with individual patients modelled as random effects. **b,c,** Spearman correlation between TNF signalling fold change and TNF signalling score pre-therapy in responders to anti-TNF treatment in **(b)** CD and **(c)** UC. DC, dendritic cell; EEC, enteroendocrine cell; GC, germinal centre; hi: high; ILC, innate lymphoid cell; lo, low; MNP, mononuclear phagocyte; Th, CD4^+^ T helper cell; Treg, CD4^+^ regulatory T cell.

**Extended Data Fig. 12.**
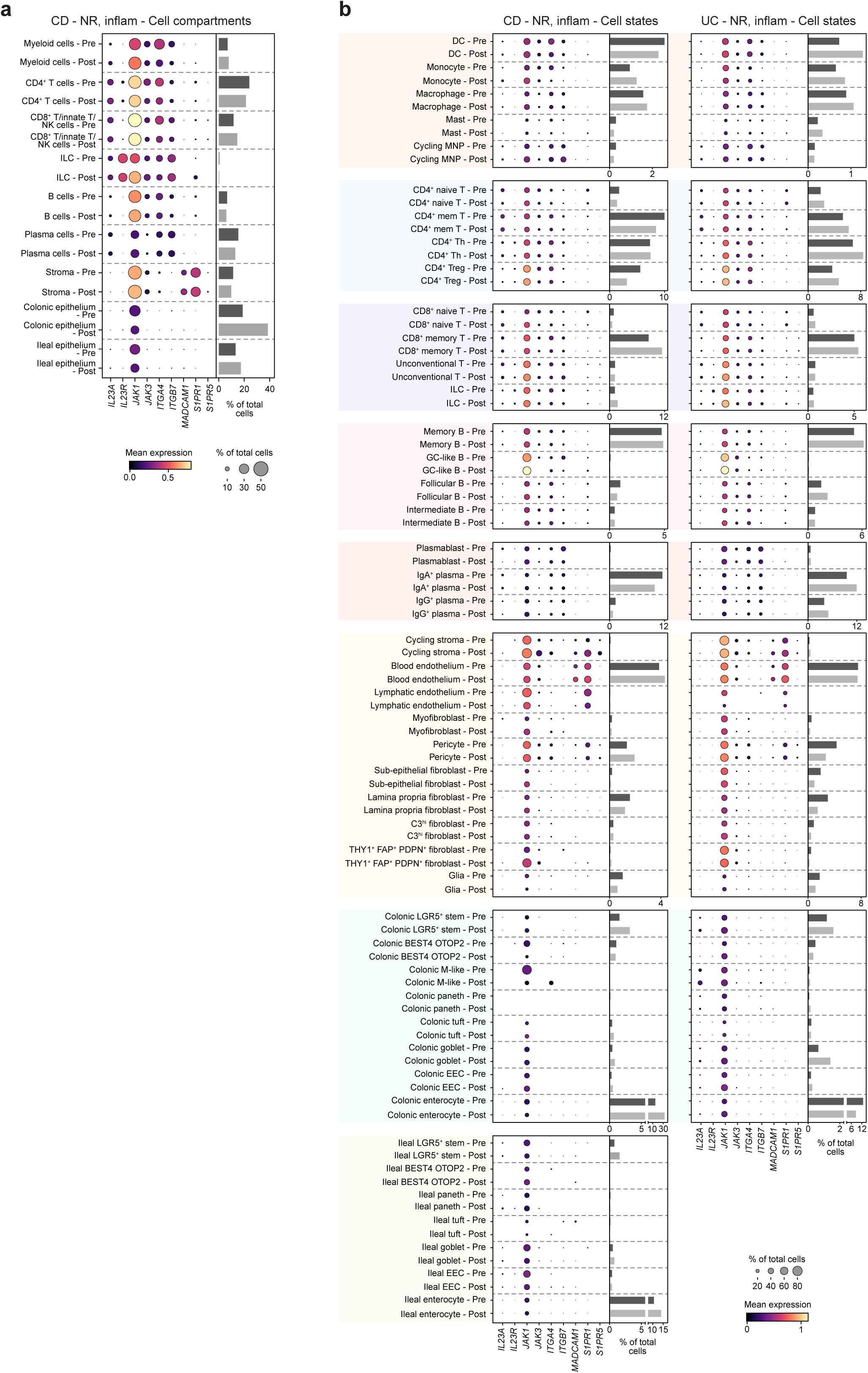
Expression of genes associated with approved advanced therapies in CD and UC in post-anti-TNF samples from non-responders. **a,** Dotplot showing expression of genes associated with advanced therapies before and after therapy at the compartment level in inflamed samples (inflam) from non-responders (NR) with CD. Bar chart shows median abundance of cell compartment in context of treatment (pre and post) as proportion of total cells in sample. **b,** Dotplot showing expression of genes associated with advanced therapies before and after anti-TNF therapy at the ‘intermediate’ level of resolution in inflamed samples (inflam) from non-responders (NR) with CD and UC. Bar chart shows median abundance of intermediate cell subpopulation in context of treatment (pre and post) as proportion of total cells in sample.

**Extended Data Fig. 13.**
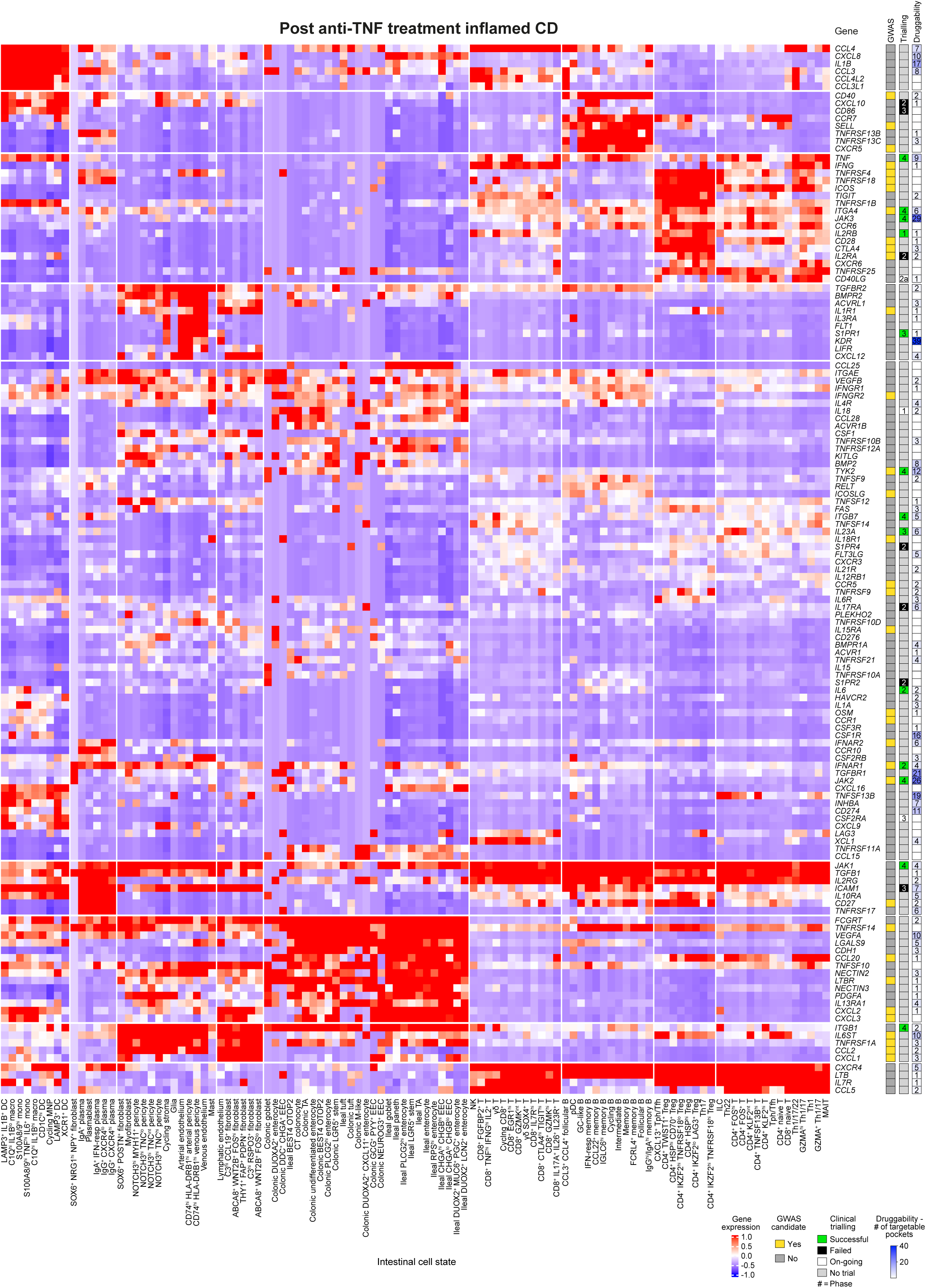
Therapeutic atlas for CD. Inflamed samples with CD following treatment with anti-TNF were pseudobulked at the cell-state resolution. A list of therapeutically relevant genes including curated cytokine and receptors from KEGG (M9809)^68^, members of the JAK family, checkpoint co-inhibitory and co-stimulatory molecules, and cell trafficking molecules was compiled. Genes with expression in over 97% of cells were kept. Column-wise, and row-wise k-means clustering was applied. The first column to the right of the genes indicates whether the gene has been implicated in genome-wide association studies (GWAS; yellow). The second column indicates stage of development of therapeutic agent associated with the gene (Phase 1/2/3/4), green colour indicative of trial success. The third column is indicative of the number of druggable pockets as outlined on Pi^69^. DC, dendritic cell; EEC, enteroendocrine cell; GC, germinal centre; hi, high; IFN-resp, interferon-responsive; ILC, innate lymphoid cell; lo, low; macro, macrophage; MAIT, mucosal-associated invariant T; MNP, mononuclear phagocyte; mono, monocyte; NK, natural killer cells; pDC, plasmacytoid dendritic cell; peri, pericyte; TA, transit-amplifying; Tfh, CD4^+^ follicular helper T cell; Tph, CD4^+^ peripheral helper T cell; Th, CD4^+^ T helper cell; Treg, CD4^+^ regulatory T cell.

**Extended Data Fig. 14.**
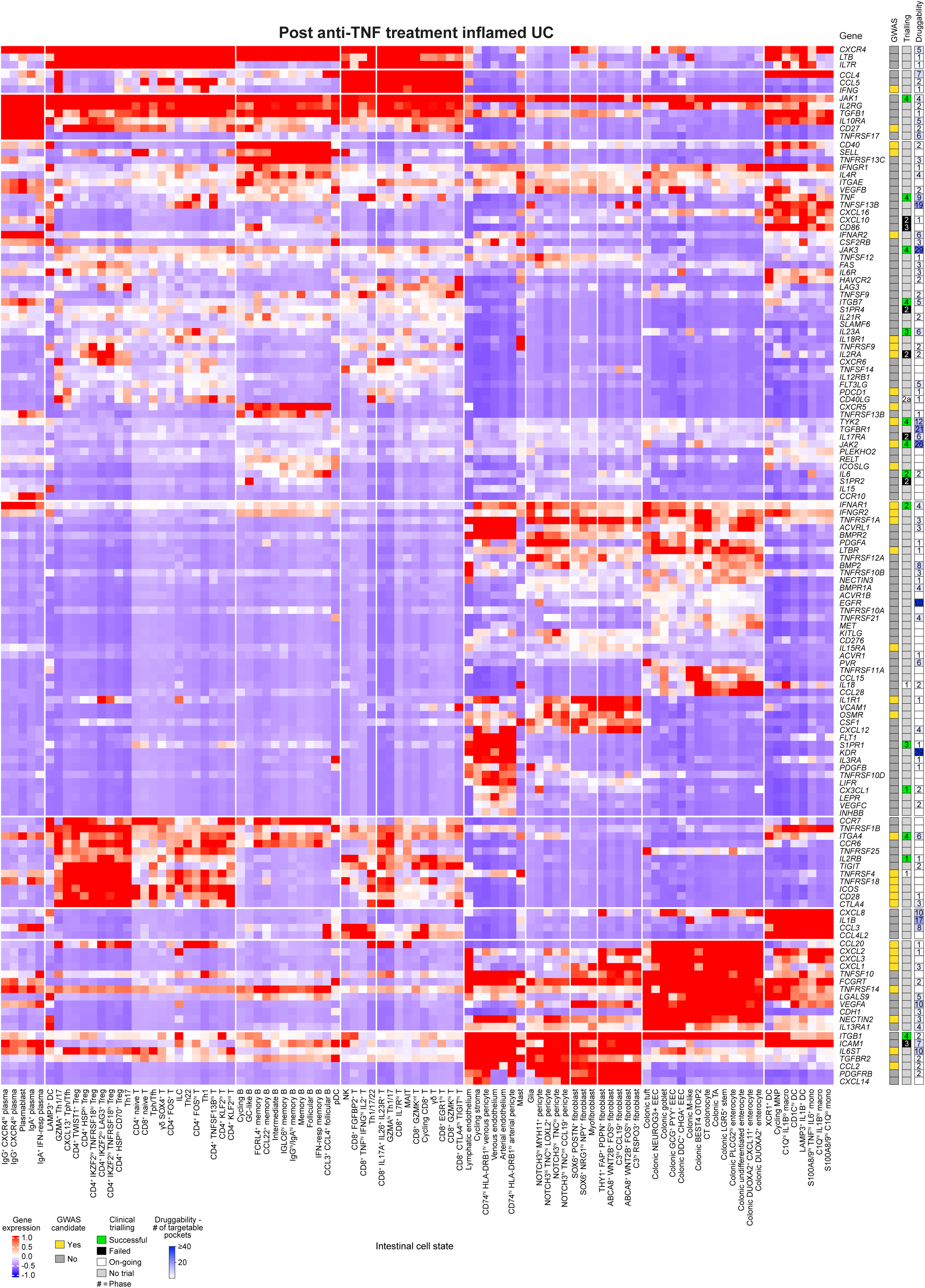
Therapeutic atlas for UC. Inflamed samples with UC following treatment with anti-TNF were pseudobulked at the cell-state resolution. A list of therapeutically relevant genes: curated cytokine and receptors from KEGG (M9809)^68^, members of the JAK family, checkpoint co-inhibitory and co-stimulatory molecules, cell trafficking molecules was compiled. Genes with expression in over 97% of cells were kept. Column-wise, and row-wise K-means clustering applied. The first column to the right of the genes indicates whether the gene has been implicated in in genome-wide association studies (GWAS; yellow). The second column indicates stage of development of therapeutic agent associated with the gene (Phase 1/2/3/4), green colour indicative of trial success. The third column is indicative of the number of druggable pockets as outlined on Pi^69^. DC, dendritic cell; EEC, enteroendocrine cell; GC, germinal centre; hi, high; IFN-resp, interferon-responsive; ILC, innate lymphoid cell; lo, low; macro, macrophage; MAIT, mucosal-associated invariant T; MNP, mononuclear phagocyte; mono, monocyte; NK, natural killer cells; pDC, plasmacytoid dendritic cell; peri, pericyte; TA, transit-amplifying; Tfh, CD4^+^ follicular helper T cell; Tph, CD4^+^ peripheral helper T cell; Th, CD4^+^ T helper cell; Treg, CD4^+^ regulatory T cell.

**Extended Data Fig. 15.**
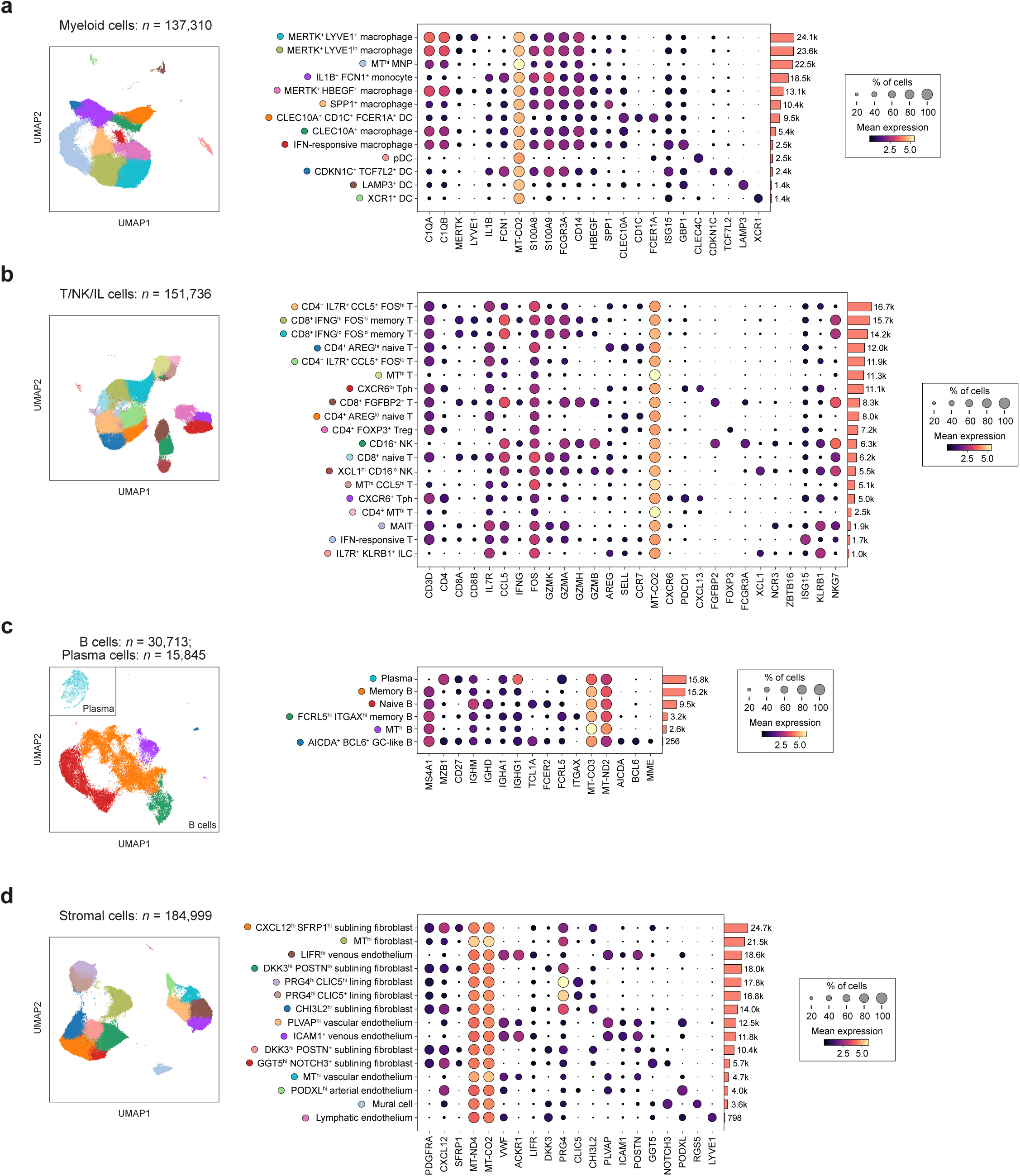
Cell states of the synovium. Uniform manifold approximation and projections (UMAPs) and associated dot plots show the expression of marker genes of cell states in the scRNA-seq dataset: **(a)** myeloid cells, **(b)** T/NK/IL cells, **(c)** B cells, **(d)** stromal cells. DC, dendritic cell; fibro, fibroblast; GC, germinal centre; hi, high; IFN-resp, interferon-responsive; ILC, innate lymphoid cell; lo, low; macro, macrophage; MAIT, mucosal-associated invariant T; MNP, mononuclear phagocyte; mono, monocyte; MT^hi^, Mitochondrial high; NK, natural killer cell; pDC, plasmacytoid dendritic cell; Tfh, CD4^+^ follicular helper T cell; Tph, CD4^+^ peripheral helper T cell; Treg, CD4^+^ regulatory T cell.

**Extended Data Fig. 16.**
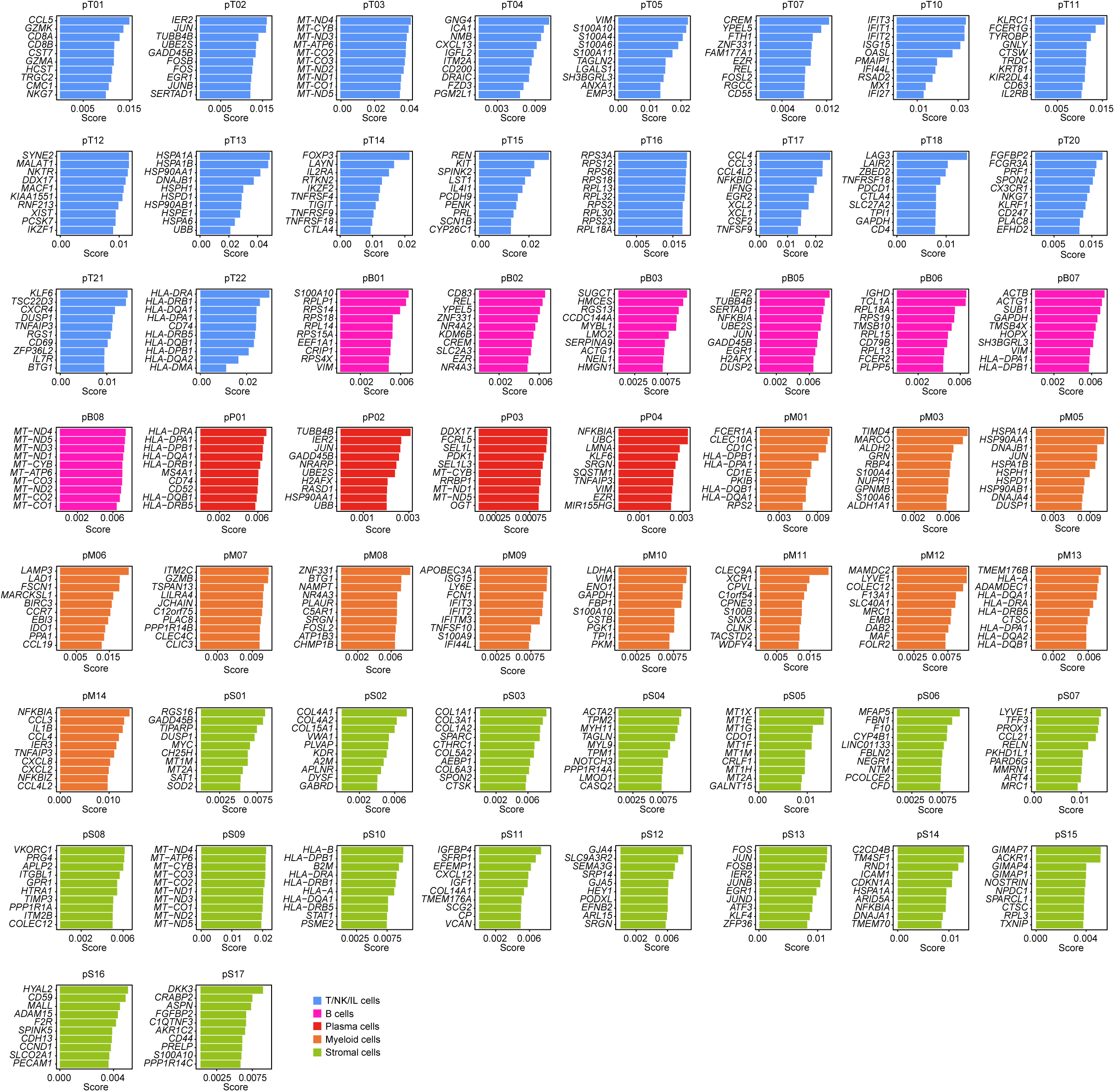
Top weighted genes of GEPs in the synovium. Weighted genes for each gene expression programme (GEP) derived from the synovium. cNMF was run separately in: T cells, B cells, plasma cells, myeloid cells and stromal cells. See **Supplementary Table 12** for full list of weighted genes, as well as results of overrepresentation analysis. See **Supplementary Table 13** for results of enrichment testing of GEPs in inflammation. pB, B cell GEP; pM, myeloid cell GEP; pP, plasma cell GEP; pS, stromal cell GEP; pT, T/NK cell GEP.

**Extended Data Fig. 17.**
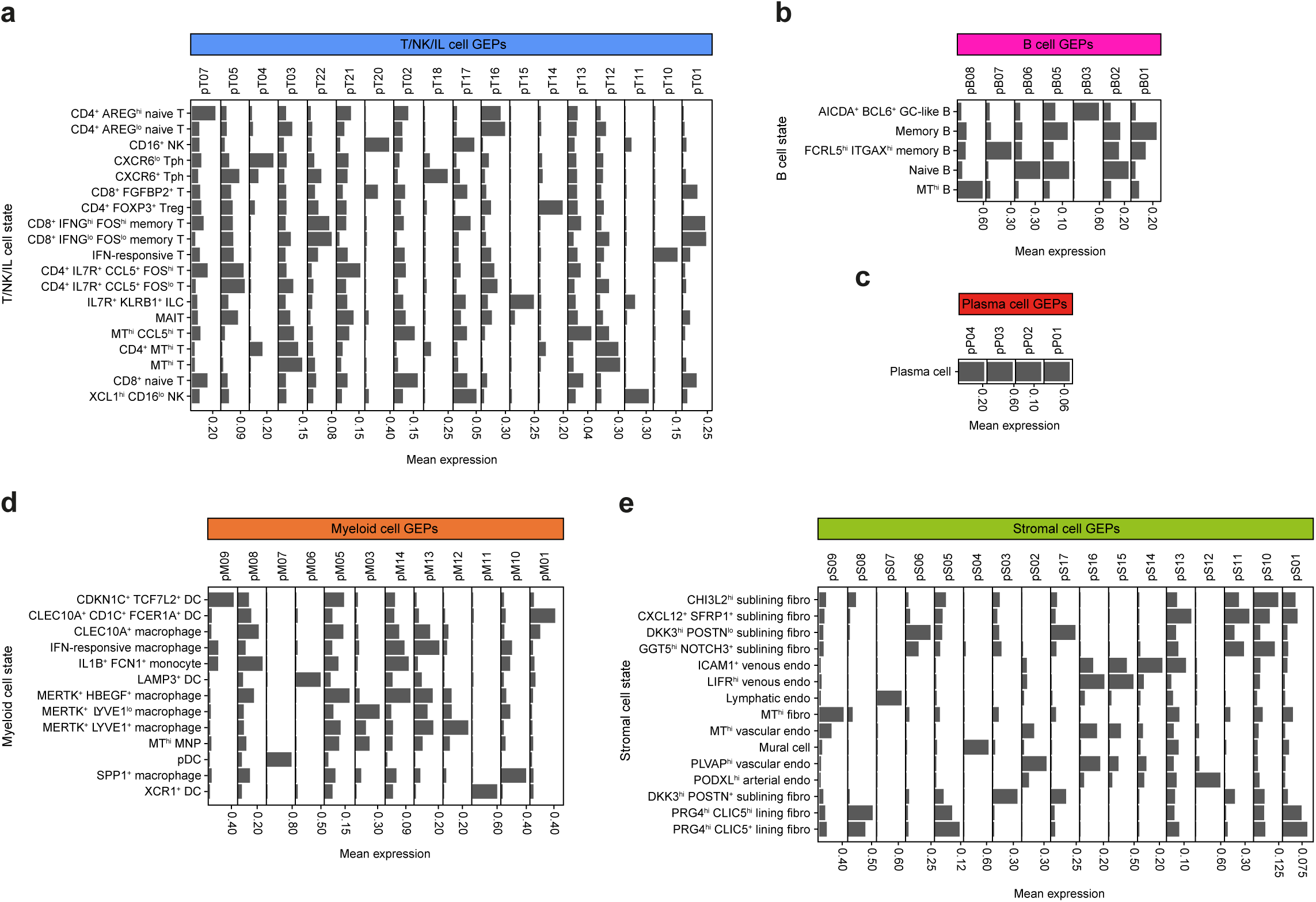
Enrichment of GEPs across cell states in the synovium. cNMF was used to derive GEP score for individual cells from inflamed samples with RA in **(a)** T/NK/IL, **(b)** B, **(c)** plasma, **(d)** myeloid, and **(e)** stromal cells. Mean expression of GEP quantified per cell state. DC, dendritic cell; endo, endothelium; fibro, fibroblast; GC, germinal centre; hi, high; IFN-resp, interferon-responsive; ILC, innate lymphoid cell; lo, low; MAIT, mucosal-associated invariant T; MNP, mononuclear phagocyte; MT^hi^: Mitochondrial high; NK, natural killer cell; pB: B cell GEP; pDC, plasmacytoid dendritic cell; pM: myeloid cell GEP; pP: plasma cell GEP; pS: stromal cell GEP; pT: T/NK cell GEP; Tfh, CD4^+^ follicular helper T cell; Tph, CD4^+^ peripheral helper T cell; Treg, CD4^+^ regulatory T cell.

**Extended Data Fig. 18.**
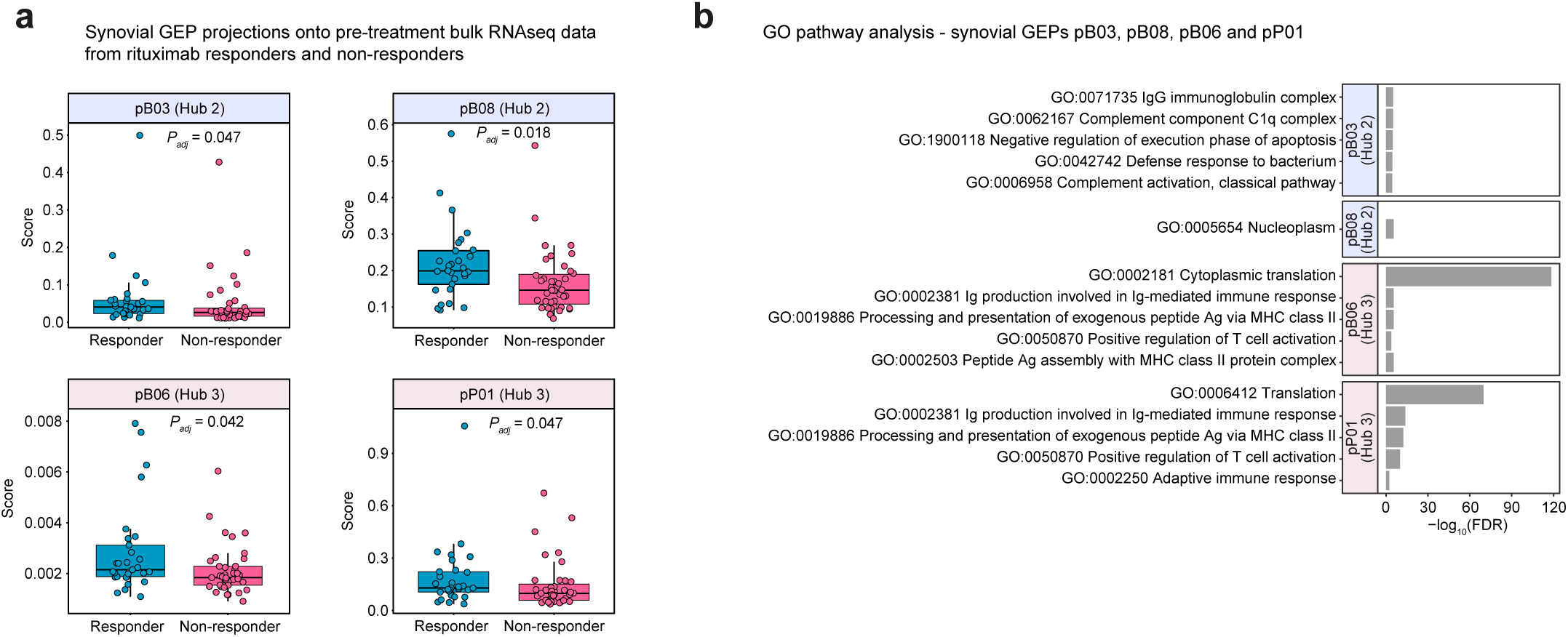
Association of synovial GEPs with clinical response to rituximab and GO pathway enrichment analysis. **a,** Baseline visit samples in the R4RA study was selected for analysis^42^. GEPs positively correlated with inflammation were tested for association with therapy non-response. Wilcoxon signed-rank test used to test for significance (*P_adj_* < 0.05) between responders (29 patients) and non-responders (39 patients) to rituximab (*n*=68 patients) at baseline. See **Supplementary Table 13** for full results. **b**, GO term enrichment for GEPs associated with clinical response to rituximab. See **Supplementary Table 13** for full results. GO terms were generated through applying overrepresentation analysis through GOATOOLS to the top 150 weighted genes in constituent GEPs^65^. All genes tested were used as the gene universe. See **Supplementary Table 12** for full list of cNMF GEPs in RA and associated GO term enrichment in GEPs. pB, B cell GEP; pP, plasma cell GEP.

**Extended Data Fig. 19.**
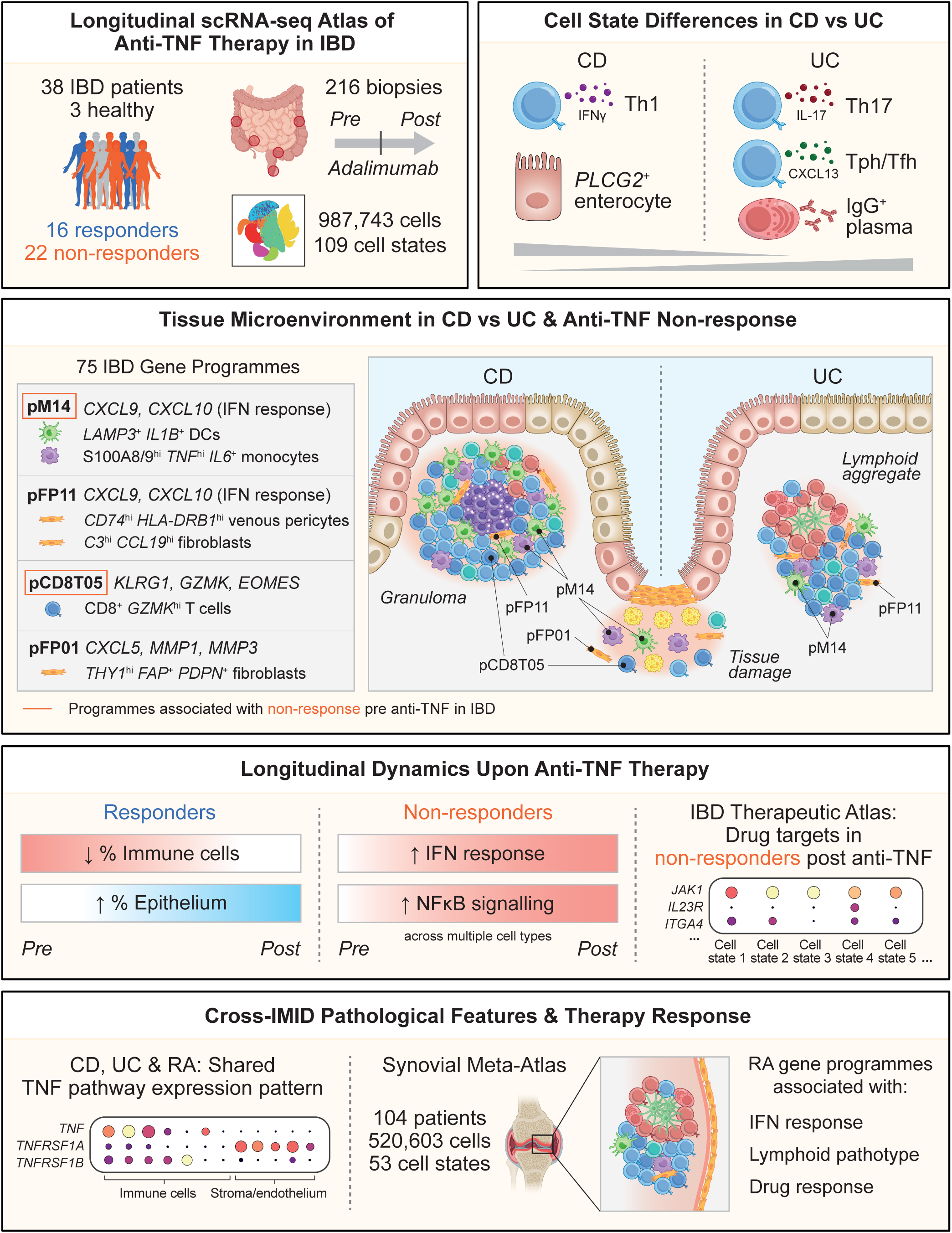
A longitudinal single-cell therapeutic atlas of anti-TNF treatment in IBD. Schematic summarising the TAURUS study design and key findings. Our resource provides a longitudinal, therapeutic scRNA-seq atlas comprising ~1 million cells organised into 109 cell states from 216 gut biopsies across 41 individuals (16 responders, 22 non-responders, 3 healthy). This atlas reveals differences in gut cell state abundance that distinguish CD and UC. Using a systems-biology approach we identify hubs of multi-cellular communities, based on 75 IBD gene programmes, which localise to distinct tissue microenvironments including granulomas specific to CD and areas of epithelial tissue damage and lymphoid aggregates found in both CD and UC. Specific programmes are associated with anti-TNF resistance pre-treatment, thereby linking drug response to spatial niches in the inflamed gut. Analysis of the longitudinal dynamics of therapeutic perturbation demonstrates that whilst cellular remission occurs in responders, in non-responders disease progression is associated with increased multi-cellular IFN and NF-kappaB signalling after anti-TNF. Extending the study to RA through the generation of a synovial meta-atlas comprising 520,603 cells reveals a shared TNF pathway expression pattern in CD, UC and RA, as well IFN signalling associated with a lymphoid pathotype. Our therapeutic atlas informs drug positioning across IMIDs, and suggests a rationale for the use of JAK inhibition following anti-TNF resistance. DC, dendritic cell; pCD8T, CD8^+^ T cell/NK GEP; pFP, fibroblast/pericyte GEP; pM: myeloid cell GEP; Th, CD4^+^ helper T cell; Tfh, CD4^+^ follicular helper T cell; Tph, CD4^+^ peripheral helper T cell.

